# Gradual Compaction of the Central Spindle Decreases its Dynamicity as Revealed in PRC1 and EB1 Gene-Edited Human Cells

**DOI:** 10.1101/2020.07.09.195347

**Authors:** Jayant Asthana, Nicholas I. Cade, Davide Normanno, Wei Ming Lim, Thomas Surrey

**Author notes:** UK Dementia Research Institute at UCL, Gower Street, London WC1E 6BT, UK.

## Abstract

During mitosis the spindle undergoes morphological and dynamic changes. It reorganizes at the onset of anaphase when the antiparallel bundler PRC1 accumulates and recruits central spindle proteins to the midzone. Little is known about how the dynamic properties of the central spindle change during its morphological changes in human cells. Using gene editing, we generated human cells that express from their endogenous locus fluorescent PRC1 and EB1 to quantify their native spindle distribution and binding/unbinding turnover. EB1 plus end tracking revealed a general slowdown of microtubule growth, while PRC1, similar to its yeast orthologue Ase1, binds increasingly strongly to compacting antiparallel microtubule overlaps. KIF4A and CLASP1 bind more dynamically to the central spindle, but also show slowing down turnover. These results show that the central spindle gradually becomes more stable during mitosis, in agreement with a recent ‘bundling, sliding and compaction’ model of antiparallel midzone bundle formation in the central spindle during late mitosis.

## INTRODUCTION

During cell division, the mitotic spindle that is responsible for segregating the two sets of chromosomes to the new daughter cells, undergoes dramatic morphological changes. During anaphase which lasts ∼ 10 min in animal cells, chromosomes are pulled to the spindle poles, while the spindle dynamically reorganises and elongates^1^. At the same time, microtubules that constitute major elements of the spindle become more stable, with their average lifetimes increasing from ∼ 20 s in metaphase to several minutes in anaphase and telophase^2-6^.

As the overall microtubule network becomes less dynamic, the spindle concentrates antiparallel microtubule overlaps in its centre, a region also called the midzone, that is critically important for spindle stability and function, as well as for intracellular signalling events ensuring correct cytokinesis^7, 8^. These antiparallel microtubule overlaps are formed and stabilised by microtubule-crosslinking proteins. PRC1 (Protein Regulator of Cytokinesis 1) is an evolutionary conserved crosslinker that connects microtubules in an antiparallel configuration^9-11^. In metaphase, PRC1 localises to “bridging fibres” which are mixed polarity microtubule bundles shown to connect opposing sister kinetochore fibres^12, 13^. As the spindle reorganises in anaphase as a consequence of the decreasing activity of Cdk1/cyclin B kinase, PRC1 is dephosphorylated and concentrates at the forming spindle midzone^14-22^. PRC1 also recruits several other midzone proteins, including motor proteins, kinases and signalling proteins^23-25^. Important examples are KIF4A, a plus-end directed kinesin that dampens microtubule dynamicity selectively in the spindle midzone ^9, 14, 21, 22^, and CLASP1/2, TOG domain containing proteins that stabilise microtubules by promoting rescues (transition from shrinkage to growth) and suppress catastrophes ^26-28^. Depletion of either KIF4A or CLASP1 results in a perturbed central spindle^21, 28^. Together all midzone proteins generate central antiparallel microtubule overlaps which have a length of ∼ 2 μm in animal cells^16, 17, 29, 30^.

In vitro experiments with purified proteins and computer simulations have provided additional insight into the molecular mechanism of antiparallel microtubule overlap formation. In vitro PRC1 is the main driver for antiparallel microtubule bundling^9-11^. KIF4A stops microtubule plus end growth selectively of overlap microtubules when it is recruited in sufficient amounts by PRC1 ^9, 31^. Additionally, together with PRC1, KIF4A slides antiparallel microtubules apart, shortening the overlaps and compacting the crosslinking proteins within the overlap which then resist further compaction leading to the formation of microtubule overlaps with a controlled length in the micrometer range, explaining why antiparallel microtubule overlaps do not slide apart completely in the late central spindle^32, 33^. In reconstituted systems, the binding/unbinding turnover of PRC1 in compacted midzone-like bundles is extremely low, whereas KIF4A is more dynamically bound ^33^. In contrast to KIF4, CLASP1 does not act selectively on overlap microtubules in vitro, because it binds considerably more weakly to PRC1 than KIF4A^34^.

Whereas the dynamic properties of the conserved PRC1 homolog Ase1 have been investigated in budding and fission yeast spindles^35-37^, much less is known about the binding/unbinding turnover of PRC1 on microtubules during different stages of mitosis in animal cells^17^. The same is true for other central spindle proteins like KIF4A and CLASP1/2. This limits our understanding of the structure and dynamic properties of the spindle during its reorganization in mitosis. Therefore, it is unclear to which extent the model for the molecular mechanism of midzone bundle formation derived from biochemical in vitro reconstitutions and computer simulations applies to the situation in the cell.

Another important cellular recruitment factor for the microtubule cytoskeleton is the end binding protein EB1^23, 38^. EB1 family members (EBs) track growing microtubule plus ends dynamically, with their N-terminal calponin homology domain binding preferentially to the growing part of microtubules where the GTP cap is^39-45^. The C-terminal part recruits binding partners that either contain a CAP-Gly domain or linear sequence motifs such as a SxIP motif ^46, 47^. ^28, 48-50^. EBs are often used as ectopically expressed fluorescent microtubule plus end markers in living cells ^17, 42, 50-64^. Overexpressed EB1 with a C-terminal fluorescent tag revealed that microtubule growth slowed down in Hela cells from metaphase to telophase by ∼ 30% ^53, 64^. However, no quantitative information is available about the plus end tracking dynamics of EB1 in the spindle under conditions avoiding overexpression and C-terminal tagging that may interfere with the interaction with binding partners^65^.

Here, we examined the distribution and the dynamic properties of PRC1 and EB1 in spindles of diploid human cells during mitosis. To allow measurements under close-to-native conditions, we engineered RPE1 cells using CRISPR/Cas9 gene-editing technology to express fluorescently tagged PRC1 and EB1 from their endogenous genetic loci. Microtubule growth speeds in the central part of the spindle slowed down from metaphase to telophase, as indicated by plus-end tracking of N-terminally fluorescently tagged EB1 at close to natural expression levels. We also quantified the redistribution of fluorescently tagged PRC1, imaging the transformation of bridging fibres in metaphase to central midzone bundles in anaphase and telophase. We found that PRC1 binding/unbinding turnover slowed down dramatically, as microtubule overlaps compacted. KIF4A and CLASP1 turnover also slowed down from anaphase to telophase although these proteins bind more dynamically to the spindle than PRC1. These results demonstrate, that central spindles become less dynamic from early anaphase to telophase, largely in agreement with quantitative models of antiparallel microtubule overlap formation and shortening in the spindle midzone.

## RESULTS

### N-terminally mGFP-tagged EB1 tracks microtubule ends in interphase and mitosis

To visualize how the microtubule network reorganizes in human cells during mitosis under close-to-native conditions, we generated hTERT-RPE1 cell lines in which PRC1 and EB1 were fluorescently tagged at their endogenous locus using CRISPR/Cas9 gene editing. We first generated a cell line with a mGFP gene inserted at the 5’-end of the EB1 gene separated by a linker to tag the N-terminus of EB1 in order not to disturb its ability to recruit other +TIPs via its C-terminus which might alter microtubule dynamics. We imaged mGFP-EB1 expressed from its endogenous locus in these gene-edited cells over the time course of mitosis in the presence of the DNA marker Hoechst. To synchronise the cells, they were arrested before mitosis at the G1/S transition by the addition of thymidine and then released by thymidine removal. Confocal microscopy imaging demonstrated that endogenously expressed EB1 with a N-terminal mGFP tag preferentially localized to growing microtubule ends both in interphase and during mitosis (Suppl. Movie 1), as previously observed for overexpressed C-terminally tagged EB1^57^ or N-terminally tagged EB3^42^. mGFP-EB1 also accumulated to spindle poles as noted previously^63^.

Using CRISPR/Cas9 gene-editing, we then inserted additionally a mCherry gene at the 5’-end of the PRC1 gene. Confocal live cell imaging revealed that in contrast to the cells in which only EB1 was tagged, in many of these doubly gene-edited cells mGFP-EB1 surprisingly colocalized together with mCherry-PRC1 at the central spindle (Suppl. Fig 1A). However, some of the doubly gene-edited cells showed the typical microtubule plus end tracking behaviour, both in interphase (Fig. 1A) and during mitosis (Fig. 1B). Several controls were performed to confirm that plus end tracking and not the central spindle localization is the native behaviour of EB1 during mitosis (Suppl. Fig. 1B-D). Using flow cytometry and confocal microscopy, we isolated single-cell clones of the doubly gene-edited cells in which EB1 displayed its normal microtubule plus-end tracking behaviour. Why EB1 can also display an atypical localization in some clones of the gene-edited cells after double knock-in, remains unknown. These atypical cell clones were not further investigated.

**Figure 1.**
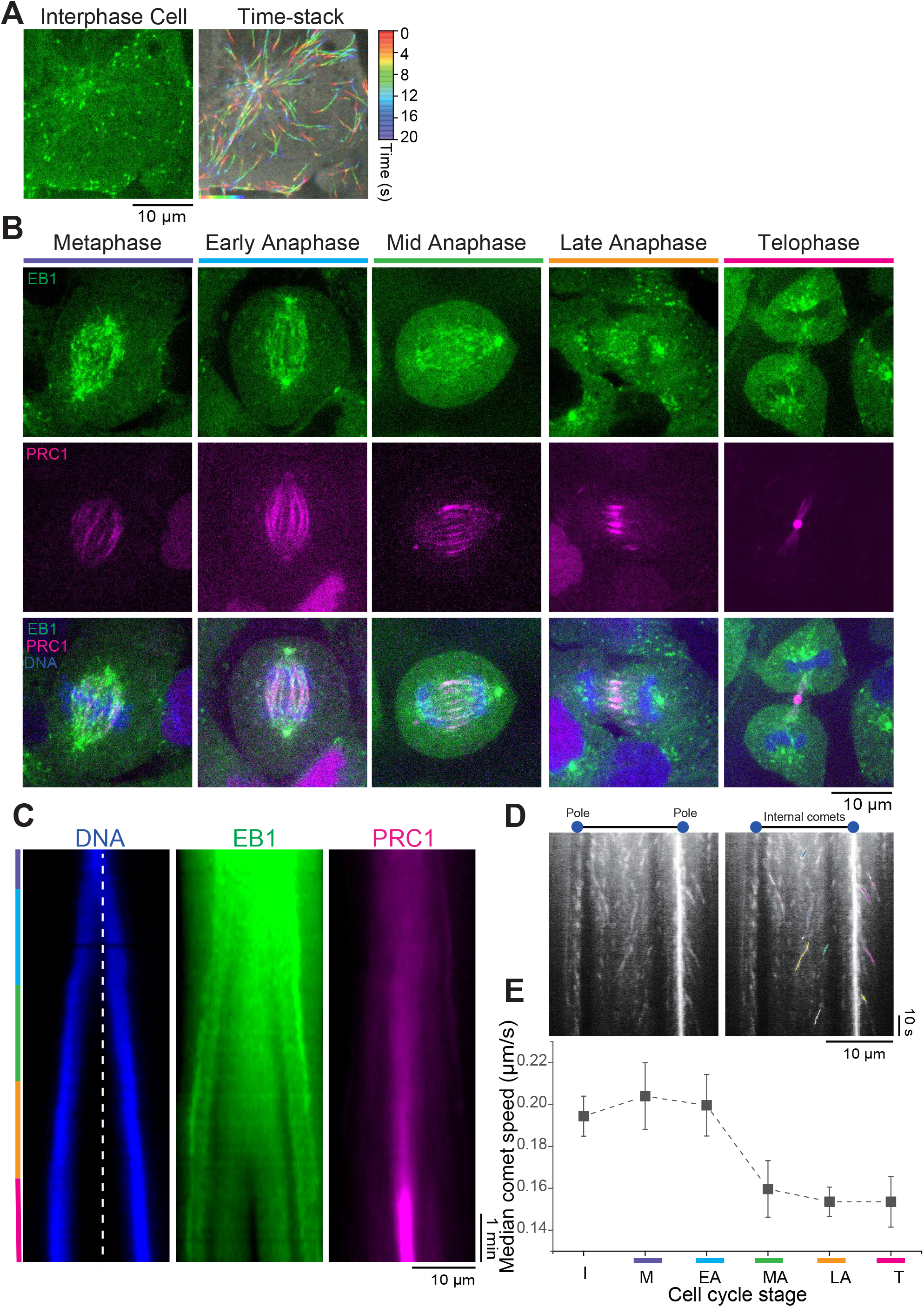
Live cell imaging of endogenously expressed mGFP-EB1 and mCherry-PRC1 during mitosis. **(A, B)** Spinning disk confocal microscopy of endogenous mGFP-EB1 in RPE1 cells (also expressing mCherry-PRC1). **(A)** Image of an interphase cell (left) and a time stack from a time lapse movie of the same cell (right) showing EB1 microtubule end tracking traces color-coded as a function of time. See also Movie S1. **(B)** Confocal microscopy images of gene-edited RPE1 cells co-expressing mGFP-EB1 (green) and mCherry-PRC1 (magenta) from their endogenous loci. Chromosomes are stained by Hoechst (blue). Images were acquired at 4 frames per second. Time is in min:s. See also Movie S2 and S3. **(C)** Representative kymographs (space-time plots) generated from a mitotic cell movie showing chromosomes (blue), EB1 (green), and PRC1 (magenta) distribution along the spindle (pole-to-pole) axis. Images were acquired every 5 seconds from metaphase to telophase. See also Movie S2. **(D)** A representative kymograph (left) generated from a late anaphase movie along the pole-to-pole axis. Internal (i.e., in between the poles) EB1 comets tracks and the EB1 tracks along astral microtubules are visible. (Right) overlay of the kymograph on the left and EB1 tracks, shown in different colors, as detected using the KymoButler deep learning software ^73^. **(E)** EB1 comet speed behaviour in interphase and from metaphase to telophase. EB1 average comet speeds were calculated from the first and last point of each trajectory. Data point are median values and error bars represent the standard error of the mean. The number of EB1 comets (n) and cells (N) considered in the analysis is: metaphase (M) n=52, N=11; early anaphase (EA) n=51, N=5; mid anaphase (MA) n=50, N=8; late anaphase (LA) n=125, N= 8; telophase (T) n=87, N=12. Scale bars = 10 µm.

Genotypic analysis of the selected single cell clones (Fig. 1A, B) showed that the mGFP and mCherry genes were inserted at the correct genomic locus (Suppl. Fig. 2A, 2D) and that both EB1 and PRC1 alleles were modified (Suppl. Fig. 2B, 2E). This means that all expressed EB1 and PRC1 molecules, including different PRC1 splice isoforms, carry a N-terminal fluorescent tag in these cells. Western blot analysis showed that mGFP-EB1 and mCherry-PRC1 were about twofold less abundant than untagged EB1 or PRC1 in control RPE1 cells (Suppl. Fig. 2C, F). One cell line simultaneously expressing mGFP-EB1 and mCherry-PRC1 was selected for further studies (Suppl. Table 1).

### EB1 microtubule end tracking slows down over the course of mitosis

EB1 preferentially localized to growing microtubule plus ends throughout the spindle as PRC1 accumulated in the midzone over the course of anaphase, as expected (Fig. 1B, C, Suppl. Movie 2, 3). At telophase, EB1 was excluded from the midzone, indicating absence of microtubule growth in this region at this time (Fig. 1B, C, Suppl. Movie 2, 3). This was also observed in a control cell line with exogenously expressed C-terminally tagged EB1-GFP and endogenously expressed mCherry-PRC1 (Suppl. Fig. 3), demonstrating that C-terminally tagged EB1 (in the presence of untagged endogenous EB1) and endogenously expressed N-terminally tagged EB1 provide similar information about microtubule dynamics in RPE1 cells.

We next wondered whether the microtubule growth speed changed during the course of mitosis in the gene-edited RPE1 cells expressing both mGFP-EB1 and mCherry-PRC1. Kymograph analysis of endogenous mGFP-EB1 (Fig. 1D, Suppl. Fig. 4) revealed that microtubules grew slightly faster during metaphase compared to interphase (Fig. 1E). As mitosis progressed, their growth speed decreased reaching a minimum at late anaphase and telophase, also in agreement with previous observations using overexpressed EB1-GFP.

### Antiparallel microtubule overlaps marked by PRC1 shorten during mitosis as the spindle elongates

We next generated a different cell line to examine more quantitatively the development of antiparallel microtubule overlaps in the central spindle during mitosis. For these experiments, we gene-edited hTERT-RPE1 cells to insert a mGFP gene at the 5’ end of the PRC1 gene to be able to image a fluorophore attached to PRC1 that is less prone to photobleaching. Genotypic analysis of single cell clones showed that the mGFP gene was inserted at the correct genomic locus (Suppl. Fig. 5A) and that both PRC1 alleles were modified (Suppl. Fig. 5B). The clone that expressed the most similar level of fluorescently tagged PRC1 compared to untagged PRC1 in control cells (Suppl. Fig. 5C) was selected for further studies (Suppl. Table 1).

We imaged mGFP-PRC1 in these gene-edited cells after synchronization with a thymidine block over the time course of mitosis in the presence of Hoechst. As soon as the nuclear envelope broke down, indicative of the beginning of prophase, PRC1 started to localize to the spindle (Fig. 2A, Suppl. Movie 4). During metaphase when chromosomes were aligned in the spindle centre, PRC1 localized evenly to microtubule bundles throughout the entire spindle, as noted previously by immunostaining PRC1 in fixed cells^15, 19, 66^ or by visualising over-expressed GFP-tagged PRC1 in bridging fibres of metaphase spindles in live cells ^12, 13^. Over the time course of anaphase when chromosomes were segregating, PRC1 became more confined to the central spindle where it selectively accumulated (Fig. 2A, Suppl. Movie 4), as observed previously using either immunostaining or overexpressing fluorescently labelled PRC1^14-19, 22, 30^. In telophase, PRC1 strongly accumulated at the midbody (Fig. 2A, Suppl. Movie 4), as expected^20, 22^.

**Figure 2.**
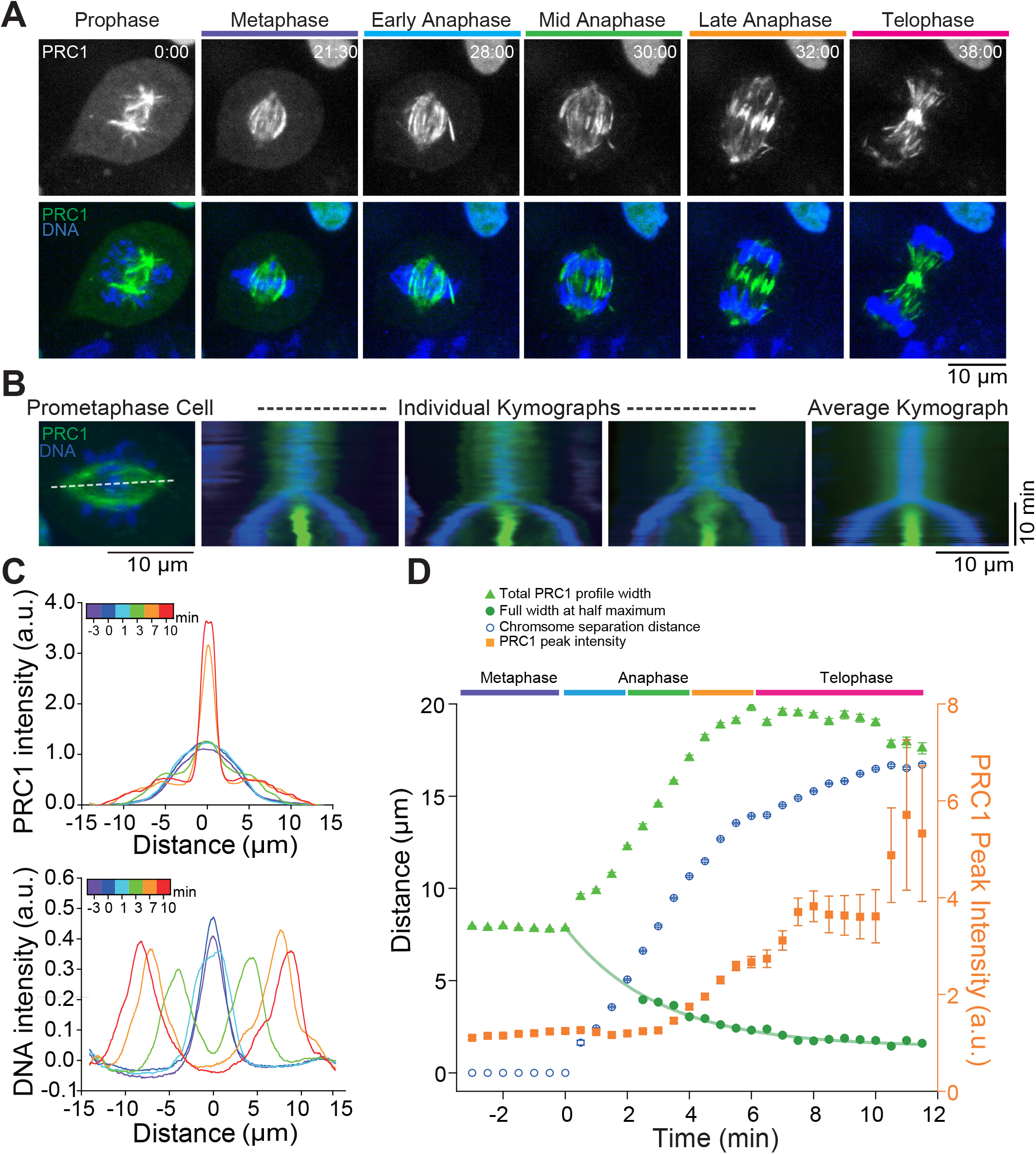
Live cell imaging of endogenously expressed mGFP-PRC1 during mitosis. **(A)** Spinning disk confocal microscopy of endogenous mGFP-PRC1 (green) and Hoechst-stained chromosomes (blue) in gene-edited RPE1 cells. Images were acquired every 30 seconds from the start of mitosis to telophase. Representative examples are shown for the different phases of mitosis as identified by the chromosome position and central spindle appearance. Scale bar = 10 µm. Time is in min:s. See also Movie S4. **(B)** Kymographs along the spindle pole-to-pole axis (white line) were generated from whole mitosis movies of RPE1 cells expressing mGFP-PRC1 (green) and Hoechst-stained to visualize the chromosomes (blue). The left image displays a pro-metaphase cell. 3 kymographs obtained from 3 different cells are shown (central images) together with the average kymograph (right image) generated from 17 individual kymographs (Methods). Scale bars = 10 µm (horizontal) and 10 min (vertical). **(C)** Example mGFP-PRC1 (top) and DNA (bottom) intensity profiles as extracted from the average kymograph. Profiles with error bars are shown in Suppl. Fig. 6B. **(D)** Time courses of total PRC1 profile width, PRC1 central peak width, and maximum intensity as a function of time (from -3.0 to 11.5 minutes after mitosis onset) as extracted from fits to the average mGFP-PRC1 intensity profile, and of the chromosome separation as extracted from fits to the average DNA intensity profile (see Methods). Green line is a fit to the PRC1 central peak width data using a mono-exponential decay function. Error bars indicate the standard error either as obtained directly from the fit to the intensity profiles or, for derived parameters, after error propagation. When not visible, errors bars fall within symbol dimensions.

Individual kymographs (space-time plots) generated along the pole-to-pole axis of the spindle and an average kymograph generated from 17 individual spindles illustrate the strong accumulation of PRC1 in the spindle centre during anaphase (Fig. 2B, Suppl. Fig. 6A). mGFP-PRC1 and DNA intensity profiles obtained from the average kymographs show that after chromosomes started to separate, the PRC1 profile transformed from a roughly Gaussian distribution in metaphase to a distribution with one central peak and two lower shoulders or side peaks (Fig. 2C, Suppl. Fig. 6B). The central PRC1 peak represents the midzone with antiparallel microtubules, whereas the side peaks likely represent spindle regions with mostly parallel microtubules to which PRC1 binds only weakly. Fitting these anaphase profiles using a sum of three Gaussians allowed us to extract the time evolution of both the overall spindle length and the central antiparallel overlap length (Fig. 2D, Methods).

During the first ∼ 5 min of anaphase, spindles in these RPE1 cells extended from an initial length of ∼ 8 μm roughly at the same rate as chromosomes separated up to a maximum length of 20 μm (Fig. 2D). After the start of cytokinetic furrow ingression (at ∼ 4 min of anaphase, not shown) the chromosomes continued to separate more slowly while spindle length did not change anymore. The central antiparallel microtubule overlaps shortened initially fast (although their exact length could not be reliably extracted from our fits until ∼ 3 min of anaphase) and then slowly approached a final length of ∼ 1.7 μm (full width at half maximum FWHM) (Fig. 2D), in agreement with previous measurements in Hela and RPE1 cells using overexpressed fluorescently tagged PRC1^17^. The density of the PRC1 molecules in the midzone increased strongly after about 3 min of anaphase as indicated by a strong increase of the central peak intensity of the PRC1 profile (Fig. 2D), as observed previously for overexpressed PRC1^16,30^. These results indicate that the PRC1 binding strength to the central midzone increases over time of anaphase. Taken together, our data demonstrate that in cells with natural PRC1 expression levels, the length of antiparallel microtubule overlaps continuously decreases towards a final end length in the micrometer range as PRC1 becomes more and more concentrated in the spindle midzone.

### PRC1 turnover decreases as the central spindle compacts

Recently the binding/unbinding turnover of ectopically expressed GFP-PRC1 in an anaphase RPE1 cell was found to be slow, with a fluorescence recovery time after photobleaching (FRAP) of several tens of seconds^17^. To determine how the dynamics of PRC1 binding to the central spindle develops over time during mitosis, we performed FRAP experiments in the spindle midzone using gene-edited RPE1 cells expressing mGFP-PRC1 from its endogenous locus at different mitotic stages (Fig. 3A, Suppl. Movie 5). Average intensity profiles of mGFP-PRC1 and Hoechst-stained DNA along the spindle axis, visualizing the extent of midzone compaction and chromosome position, respectively (Fig. 3B, top), together with the corresponding average mGFP-PRC1 fluorescence recovery curves at the different stages of mitosis were generated from at least 15 individual bleached spindles per condition (Fig. 3B, bottom). We observed that the recovery time increased continuously during the course of mitosis, with values of 6 s, 11 s, 11 s, 44 s and 124 s measured in metaphase, early anaphase, mid-anaphase, late anaphase and telophase, respectively (Fig. 3C, Suppl. Movie 5). The recovery fraction decreased from 0.55 to 0.27 in metaphase and telophase, respectively. A similar trend was observed previously for the PRC1 orthologue Ase1 in budding yeast spindles^35^, suggesting evolutionary conservation of increasingly strong PRC1 binding over the time of mitosis. These results demonstrate that the binding/unbinding turnover of PRC1 slows down the more the central spindle compacts, until hardly any PRC1 turnover is measurable at the midbody in telophase (Fig. 3D).

**Figure 3.**
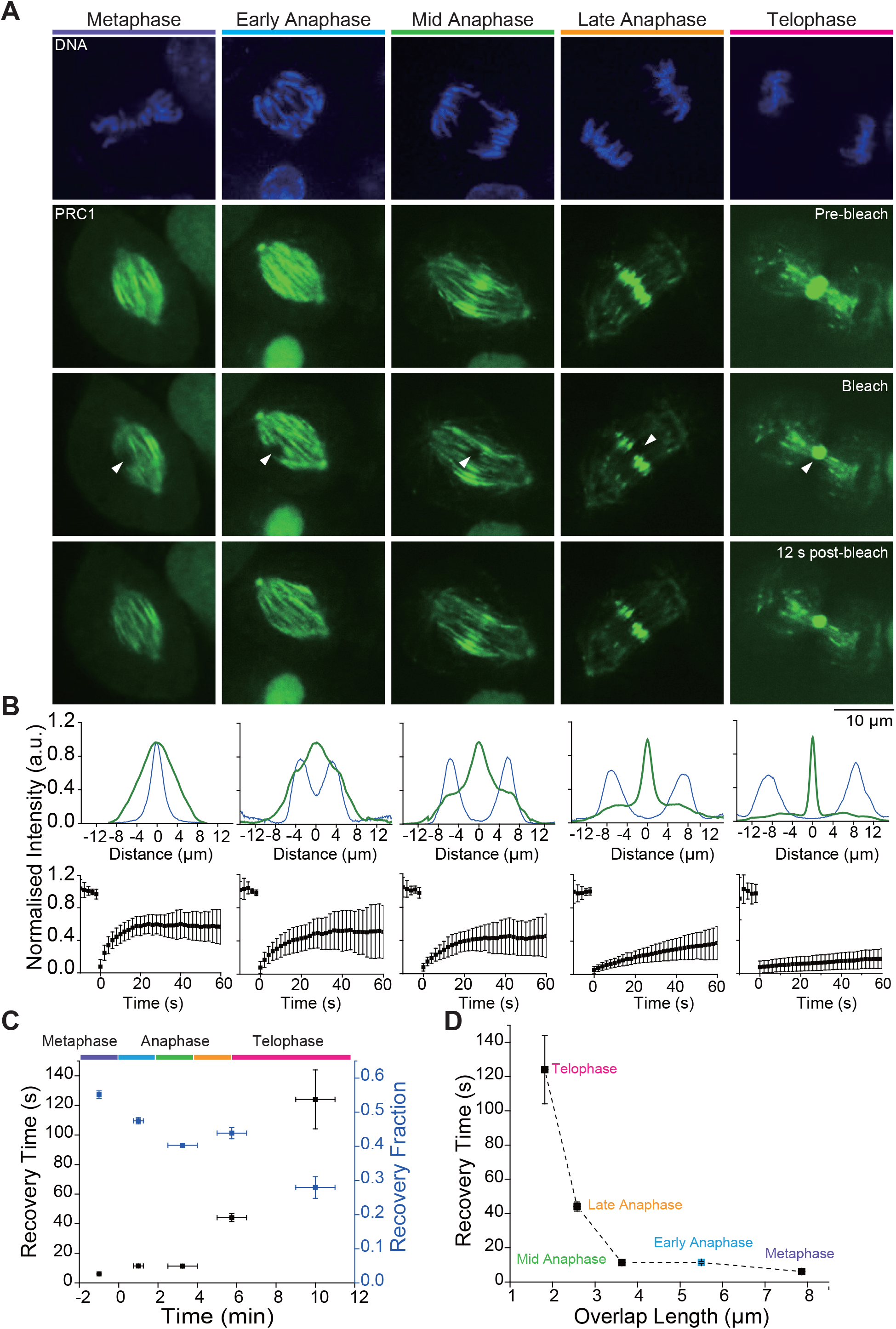
FRAP analysis of PRC1 turnover during mitosis. **(A)** Analysis of mGFP-PRC1 binding/unbinding turnover in the central spindle of gene-edited RPE1 cells at different stages of mitosis. The position of Hoechst-stained chromosomes (top row) and the binding pattern of PRC1 to the spindle (second row) were used to identify the mitotic stage. For each stage, mGFP-PRC1 bound to the spindle is shown just before photobleaching (pre-bleach, second row), at the time of bleaching (arrowheads show the bleached circular, ∼0.6 µm-radius, region; bleach, third row), and 12 s after the bleach mark was set (12 s post-bleach, bottom row). Scale bar = 10 µm. See also Movie S5. **(B)** Normalized average PRC1 (dark green) and chromosome (dark blue) intensity profiles along the spindle pole-to-pole axis at the time just before photo-bleaching (top). Average fluorescence recovery after photobleaching curves (bottom, black symbols are mean intensities, n ≥ 12 for each condition) at the different stages of mitosis shown in (A). Error bars indicate the standard deviation. **(C)**. The recovery time and the recovery fraction of mGFP-PRC1 after photobleaching at different times during mitosis. Error bars are standard errors. **(D)** The recovery time of mGFP-PRC1 after photobleaching plotted as a function of overlap length in different phases of mitosis. The overlap lengths are averages of individual overlap lengths (FWHM) from Fig. 2D for the different phases of mitosis; for early anaphase (blue symbol), the overlap length is estimated from the fit to the FWHM values in Fig. 2D. Error bars are standard errors. When not visible, errors bars fall within symbol dimensions.

### KIF4A turnover slows down in late phases of mitosis

To get a more comprehensive view of the dynamics of midzone bundles, we next asked how the turnover of central spindle proteins that are recruited by PRC1 compares to the turnover of PRC1 itself. Using lentivirus-mediated infection, cells stably expressing ectopically central spindle protein KIF4A-mGFP were generated from cells already expressing endogenous mCherry-PRC1. Confocal live cell imaging of synchronized cells revealed that in these RPE1 cells KIF4A localized to the central spindle approximately 4 minutes after anaphase onset (Fig 4A), in agreement with previous observations in HeLa cells^14^. To determine the binding/unbinding turnover to/from the central spindle of KIF4A, FRAP experiments were carried out in midzone bundles labelled by mCherry-PRC1 (Fig. 4B, C, Suppl. Movie 6). Fluorescence recovery of KIF4A showed that its turnover slowed down from anaphase to telophase with values for the recovery time of 5.9 s in anaphase and 30 s in telophase (Fig. 4D). Concomitantly, the recovery fraction decreased from 0.95 to 0.71, respectively (Fig. 4D). These values are in a similar range as a previously reported value for anaphase^67^. These data show, that KIF4A also becomes more stably bound to midzone bundles as they compact, but its binding/unbinding turnover is about one order of magnitude faster than PRC1 (Fig. 3C, D and Fig. 4C, D).

**Figure 4.**
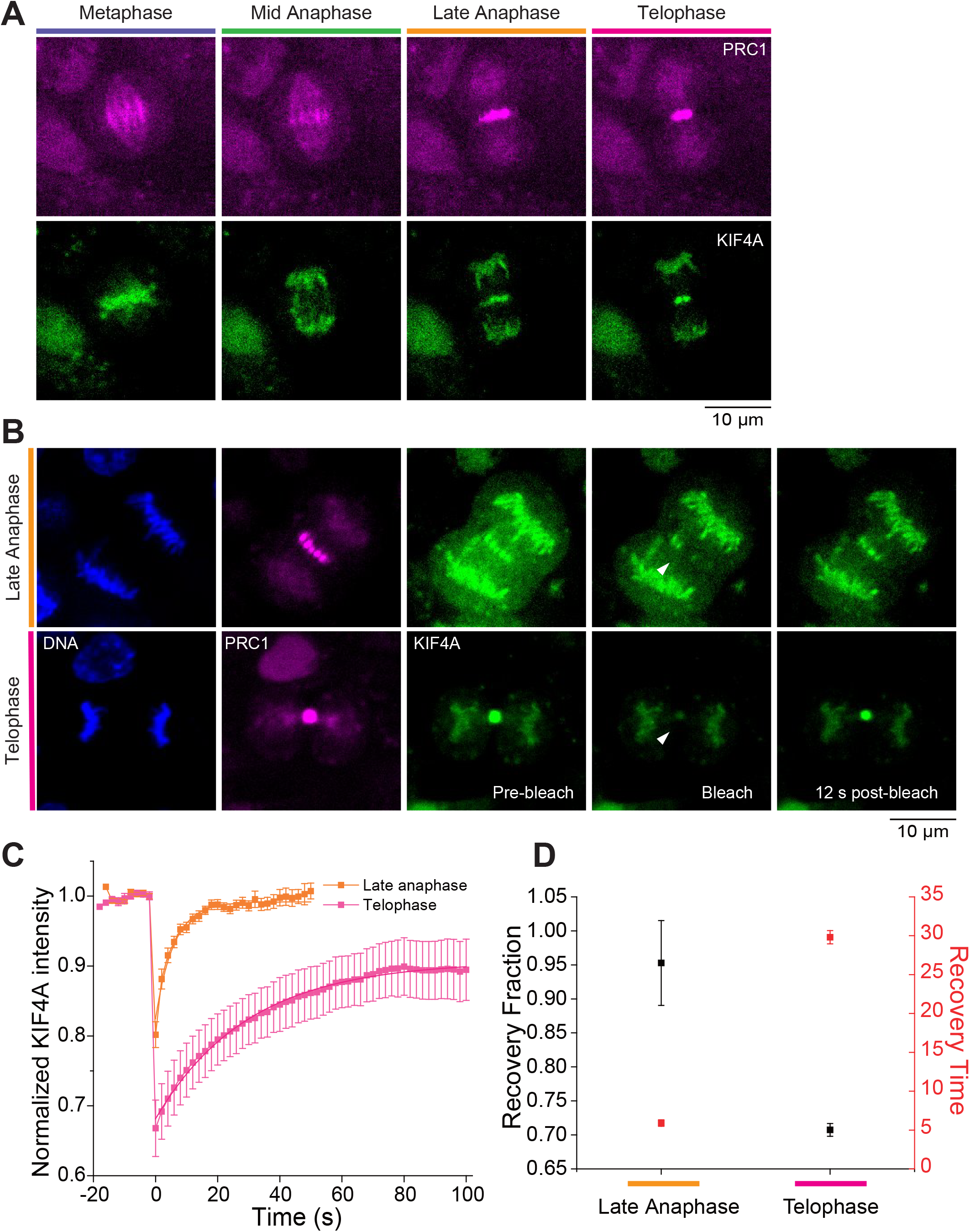
FRAP analysis of KIF4 turnover during late anaphase and telophase. **(A)** Distribution of PRC1 (magenta) and KIF4A (green) during different phases of mitosis. PRC1 is located throughout the spindle in metaphase, whereas KIF4A is on the chromosomes. KIF4A first localizes to the central spindle during late anaphase. **(B)** Analysis of KIF4A-mGFP turnover in the central spindle of RPE1 cells stably expressing ectopic KIF4A-mGFP (green) and endogenous mCherry-PRC1 (magenta) in late anaphase and telophase. The position of Hoechst-stained chromosomes (1st panel) and the binding of PRC1 to the spindle (2nd panel) are shown for late anaphase and telophase. For each stage, KIF4A bound to the spindle is shown just before photobleaching (pre-bleach, third panel), at the time of bleaching a circular region with ∼ 0.6 µm-radius (bleach, fourth panel) and 12 s after the bleach mark was set (12 s post-bleach, fifth panel). The position of photo-bleached regions is indicated by white arrowheads. Scale bar = 10 µm. See also Movie S6 **(C)** Fluorescence recovery after photobleaching curves of KIF4A in late anaphase and telophase. Data points indicate mean intensity values after normalization of the bleached regions of N=14 cells for Late anaphase, and N=18 cells for Telophase. Error bars represent the standard error of the mean. **(D)** The recovery fraction and the recovery time of KIF4A after photobleaching in late anaphase and telophase. Error bars are standard errors. When not visible, errors bars fall within symbol dimensions.

### CLASP1 turns over faster in the central spindle than both PRC1 and KIF4A

Using lentivirus-mediated infection, from cells expressing endogenous mCherry-PRC1, we generated cells also stably expressing ectopically EGFP-tagged central spindle protein CLASP1. Confocal live cell imaging of synchronized cells showed the typical localization of EGFP-CLASP1 in midzone bundles from early anaphase to telophase (Fig 5A). FRAP experiments of EGFP-CLASP1 in midzone bundles as labelled by mCherry-PRC1 revealed an even faster binding/unbinding turnover of EGFP-CLASP1in the range from 1.3 to 2.9 s (Fig. 5B) with a recovery fraction in the range from 0.64 to 0.79 (Fig. 5C). Comparing the turnover of all proteins investigated here, the turnover rate varies between proteins with CLASP1 being the most weakly bound protein and PRC1 being the protein bound most strongly to midzone bundles (Fig. 5D). Despite these differences between different central spindle proteins, the FRAP data reveal a general slowdown of protein binding/unbinding turnover in the central spindle as midzone bundles compact during mitosis.

**Figure 5.**
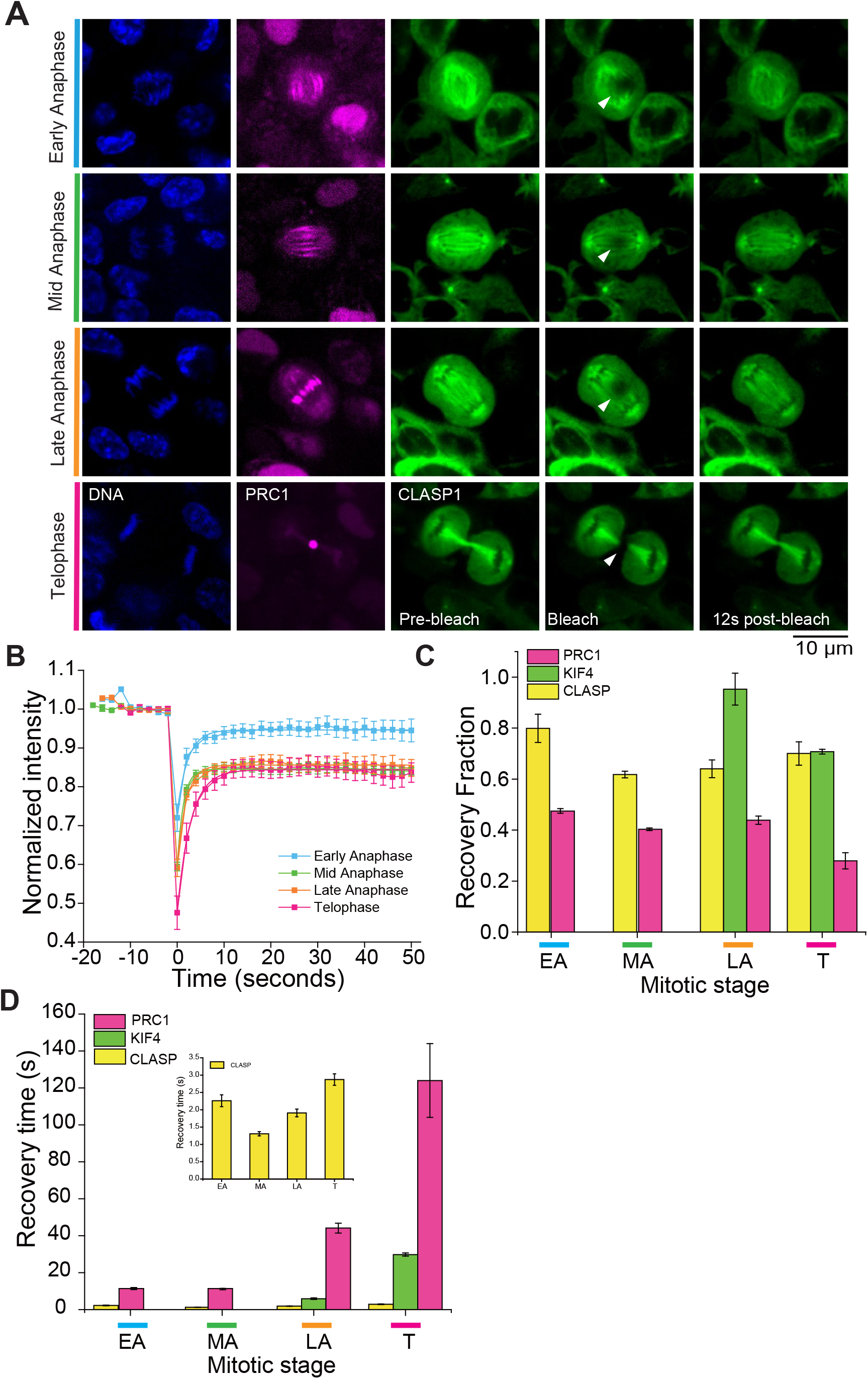
CLASP1 has a faster turnover than PRC1 and KIF4A. **(A)** Analysis of the CLASP1 turnover in the central spindle of RPE1 cells expressing ectopic EGFP-CLASP1 (green) and mCherry-PRC1 (magenta) from its endogenous loci. The position of Hoechst-stained chromosomes (1st panel) and the binding pattern of PRC1 to the spindle (2nd panel) were used to classify the mitotic stage. For each mitotic stage, EGFP-CLASP1 bound to the spindle is shown just before photobleaching (pre-bleach, third panel), at the time of bleaching a circular region with a ∼0.6 µm-radius (bleach, fourth panel) and 12 s after the bleach mark was set (12 s post-bleach, fifth panel). The position of photobleached regions is indicated by white arrowheads. **(B)** Fluorescence recovery after photobleaching curves of EGFP-CLASP1 in different phases of mitosis. Data points indicate mean intensity values after normalization of the bleached regions of N=9 cells for Early anaphase, N=10 cells for Mid anaphase, N=11 cells for Late anaphase, and N=8 cells for Telophase. Error bars represent the standard error of the mean. **(C)** Comparison of recovery fractions of PRC1 (magenta), KIF4A (green), and CLASP1 (yellow) at different mitosis stages. Error bars are standard error. **(D)** Comparison of recovery times of PRC1 (magenta), KIF4A (green), and CLASP1 (yellow) at different mitosis stages. Inset: zoom-in of the recovery time of CLASP1 at different mitosis stages. Error bars are standard error.

## DISCUSSION

We investigated how dynamic properties of the spindle change in gene-edited human RPE1 cells during mitosis. Because EB1 and PRC1 are key hub proteins recruiting a multitude of other proteins we expressed fluorescent fusion proteins of EB1 and PRC1 from their endogenous loci in an attempt to maintain at close as possible their natural expression levels.

In contrast to previous work with EB1, we tagged it at its N-terminus to preserve its ability to interact with binding partners that interact with its C-terminus, so that a situation as normal as possible could be maintained. N-terminally tagged EB1 tracked microtubule plus ends normally, revealing an overall slowdown of microtubule growth, as observed previously with overexpressed, C-terminally tagged EB1^53, 64^, possibly indicative of a reducing activity of a microtubule polymerase in response to mitosis progression.

Quantifying PRC1 fluorescence showed that antiparallel midzone bundles in the central spindle shorten from metaphase to telophase approaching a final defined length. The binding/unbinding turnover of PRC1 strongly slows down over time of mitosis, indicative of a very stable structure forming at the central spindle. KIF4A and CLASP1 are more dynamically bound to midzone bundles and differences in their turnover rates can be explained by their different affinities for PRC1 binding ^34^.

Confirming previous observations either with immuno-stained fixed cells or with live cells in which fluorescently tagged PRC1 was overexpressed, we observed that PRC1 localized to extended microtubule bundles in the metaphase spindle, where it was recently shown to localize to microtubule bundles bridging sister kinetochore fibres^12, 13, 15, 19, 66^. From anaphase on, it strongly accumulated in the spindle midzone, as previously observed^14-19, 22, 30^. Since PRC1 is known to preferentially bind antiparallel microtubules^9-11^, our data suggest that antiparallel microtubules exist throughout most of the metaphase spindle that then become focused in anaphase within ∼ 5 min into short bundled antiparallel overlaps of ∼ 1.7 μm length. The shape changes of the PRC1 profiles measured along the spindle axis over time reveal a process of longitudinal compaction, probably by a combination of motor-mediated microtubule sliding and microtubule stabilization in the midzone, and a pronounced reduction of binding/unbinding turnover of PRC1 over time.

The increasingly tighter binding of PRC1 over the time course of mitosis is very similar to what was observed for the PRC1 orthologue Ase1 in budding and fission yeast^35-37, 68^, suggesting an evolutionarily conserved stabilization of the central spindle in late mitosis. The increased binding strength of PRC1 is thought to be due to an enhanced microtubule binding affinity when PRC1 becomes dephosphorylated at the transition from metaphase to anaphase ^14, 15, 18, 19, 21, 66^. Additionally, the slowdown of its binding/unbinding turnover could be due to steric protein trapping in compacting three-dimensional antiparallel overlaps, consisting of multiple microtubules, or a combination of both factors.

Recent in vitro reconstitution experiments with purified PRC1 and the kinesin KIF4A produced microtubule bundles with antiparallel overlaps whose behaviour could be explained by computer simulations by a ‘bundle, slide and compaction’ mechanism that had similar properties as the midzone bundles characterized here in living cells ^33^. The kinetics of formation, the antiparallel overlap length and the low turnover of PRC1 and the faster turnover of KIF4A in reconstituted minimal midzone bundles mimicked the physiological behaviour measured here, despite the absence of other prominent midzone proteins. This suggests that the activities present in the in vitro system, namely antiparallel microtubule bundling, antiparallel motor sliding, and protein compaction are key activities required for the formation of a basic midzone overlap architecture.

The cell lines produced here promise to be useful tools for future studies examining quantitatively the regulatory mechanisms controlling microtubule dynamics and spindle organization over the course of mitosis.

## METHODS

### Donor vector constructs, guide RNA design and cloning

To generate human telomerase reverse transcriptase-immortalized retinal pigmented epithelium (hTERT-RPE1) cell lines expressing fluorescently tagged PRC1 and/or EB1 from their endogenous locus, we used CRISPR/Cas9-mediated gene editing to fuse mGFP or mCherry sequences to the PRC1 and/or EB1 genes^69, 70^. Homology arms of the donor vectors were made from genomic DNA of hTERT-RPE1 cells by PCR amplification of the flanking regions (1.2 kb each) of the PRC1 and EB1 genes. The coding sequences of mGFP and mCherry were PCR amplified from plasmids. Then the homology arms and fluorescent protein sequences were assembled and cloned into the BamHI site of a pUC19 vector using Gibson assembly (Clontech). The primer sequences for the mGFP-EB1, mGFP-PRC1 and mCherry-PRC1 donor constructs are listed in Suppl. Material 1 and 2. The reverse primers for the generation of the donor constructs encoded also a sequence encoding a five-glycine linker sequence to be inserted between mGFP or mCherry and PRC1 or the amino acid sequence AQAGGSGGAGSGGEGAVDG to be inserted between mGFP and EB1 (Suppl. Material 1 and 2). The latter linker is identical to the previously used linker in a GFP-EB3 construct that was overexpressed in human cells^42^. The protospacer adjacent motifs (PAM) in the donor plasmids were modified to silent mutations by site directed mutagenesis to ensure that the donor plasmids are not cleaved by Cas9 nuclease.

Guide RNA (gRNA) sequences specifying the site of Cas9 cleavage in endogenous DNA were designed from the genomic EB1 and PRC1 sequences using the guide RNA design resource (http://crispr.mit.edu/) selecting the top gRNA hits (Suppl. Material 1 and 2). Each guide RNA pair was cloned into either the ‘All-in-one-mCherry’ plasmid (AIO-mCherry, gift from Steve Jackson (Addgene plasmid # 74120; http://n2t.net/addgene:74120; RRID: Addgene_74120)) or into ‘All-in-one-GFP’ plasmid (AIO-GFP, gift from Steve Jackson (Addgene plasmid # 74119 http://n2t.net/addgene:74119; RRID: Addgene_74119))^69^. The AIO plasmids also expresses Cas9 nickase.

### CRISPR/Cas9-mediated gene editing

hTERT-RPE1 cells were cultured at 37°C and 10% CO_2_ in Dulbecco’s Modified Eagle Medium (DMEM) supplemented with 10% fetal bovine serum, 1 x non-essential amino acids (Thermofisher Scientific) and 1x penicillin-streptomycin (Thermofisher Scientific). hTERT-RPE1 cells were transfected with donor plasmid and AIO plasmid with the appropriate gRNAs using a Neon transfection system (Thermofisher Scientific). For 5 × 10^6^ cells, 5 µg of donor plasmid and 5 µg of gRNA plasmids were used for transfection (1050 V pulse voltage, 35 ms pulse width, and pulse number 2). After 16-18 hours, cells were selected by flow cytometry on the basis of expression of the fluorescence protein marker of the AIO plasmid. The selected cells were pooled and after two weeks of cell growth, flow cytometry was performed again, this time based on fluorescent protein expression of the edited endogenous genes, to collect single cells in 96 well plates. After 2 - 3 weeks when the cells became confluent, the monoclonal cultures were transferred to 24 well plates. A minimum of 30 clones per cell line to be generated were selected at this stage for further analysis. Genomic DNA was extracted and a genotypic characterization of the single cell clones was performed to verify the correct locus insertion and to determine the number of modified alleles. Primers pairs (Suppl. Material 1 and 2) consisting of one primer that binds outside of one of the homology arms in the genomic region and another primer that binds inside either the mGFP or mCherry sequence were used to verify insertion into the correct locus. Primer pairs (Suppl. Material 1 and 2) consisting of one primer that binds in the genomic region just outside of one of the homology arms and the other primer that binds inside the other homology arm, were used to detect the number of modified alleles. One hTERT-RPE1 cell clone expressing mGFP-EB1 was further edited to express also endogenous mCherry-PRC1. A pool of mGFP-EB1 and mCherry-PRC1 co-expressing cells were selected using flow cytometry. Fluorescence microscopy revealed that EB1 colocalized with PRC1 at the central spindle in 80-90% of the cells in this cell pool (Suppl. Fig. 1A), even when genotypic characterization of single cell clones demonstrated that both fluorescent protein-coding genes were inserted correctly in the genome. This pronounced PRC1 co-localization of EB1 was considered unnatural after examining untagged EB1 in control cells by immunofluorescence microscopy (Suppl. Fig. 1B-D). However, two single cell clones with correct insertions as shown by genotypic analysis showed natural EB1 behaviour as observed by fluorescence microscopy. Protein expression levels of these clones were analysed by western blotting and one of these clones was selected for further studies.

### Lentivirus-mediated stable cell line development

The EB1-GFP lentivirus expression construct was made by PCR amplification of EB1-GFP from pEGFP N1 EB1-GFP (JB131) (gift from Tim Mitchison & Jennifer Tirnauer (Addgene plasmid # 39299; http://n2t.net/addgene:39299; RRID: Addgene_39299) followed by cloning into a pLVX-puro vector (Clontech) using the XhoI and BamHI restriction sites (primers in Suppl. Material 2). EB1-GFP expressing lentivirus particles were generated by transfecting *psPAX2, pMD2*.*G* (helper plasmids), and pLVX-puro-EB1-GFP plasmids in 293FT cells. hTERT-RPE1 cells already expressing endogenous mCherry-PRC1 were transduced using the EB1-GFP lentivirus particles. hTERT-RPE1 cells stably expressing EB1 with a C-terminal GFP fusion from a randomly inserted ectopic EB1-GFP gene under the control of a CMV promoter were selected on puromycin (10 µg/ml) for 3 - 4 days. Cells expressing low levels of EB1-GFP were selected by flow cytometry and were used for live cell imaging. The expression levels were determined by western blot using EB1 monoclonal antibodies. The htert-RPE1 cells expressing endogenous mCherry-PRC1 were further modified using a lentivirus vector to express either KIF4A-mGFP or CLASP1α-EGFP. The KIF4A-mGFP lentivirus expression construct was made by PCR amplification of KIF4A-mGFP from pFastBac1-KIF4A-mGFP-His plasmid^33^ followed by cloning into pLVX-puro vector (Clonetech) using EcoRI and BamHI restriction sites (primers in Suppl. Material 2). The EGFP-CLASP1α lentivirus expression construct was made by PCR amplification of EGFP-CLASP1α from pEGFP-CLASP1α C1 plasmid^49, 71^ (a gift from Anna Akhmanova, Utrecht University, The Netherlands) and then cloned into pLVX-puro vector (Clonetech) using EcoRI and BamHI restriction sites (primers in Suppl. Material 2). The lentivirus particle generation and transduction were done as described above for EB1-GFP. The cells expressing either KIF4A-mGFP or CLASP1α-EGFP were further selected on puromycin (10 µg/ml) for 3 - 4 days. Flow cytometry was done to select the cells expressing low levels of tagged proteins.

### Live cell imaging

Cells were seeded at a density of 0.5 × 10^5^ cells/ml in 6 well plate and six hours later thymidine was added at a final concentration of 2.5 mM to block the cells at the G1/S transition. After 16-18 hours incubation, cells were washed several times with 1x PBS and once with the culture medium. After washing, cells were detached using trypsin-EDTA and seeded with fresh culture medium in 2-well dishes (Ibidi, Cat.No:80287) and kept in the CO_2_ incubator at 37°C for 8-10 hours to allow cells to release from at the G1/S transition and enter mitosis. Before imaging, the DMEM medium used to maintain the cells was replaced with DMEM without phenol red, and supplemented with 25 mM HEPES (pH 7.6) buffer. For DNA visualization, the live cell dye Hoechst 33342 was added. Mitotic cells were imaged at 37°C using a custom assembled setup (3i UK) based on a Zeiss Axio Observer Z1 equipped with a Yokogawa CSU M1 spinning disk unit, a FRAP module (Vector, 3i) and a Prime 95B sCMOS camera (Teledyne Photometrics, USA). Time lapse imaging over the entire duration of mitosis was performed in three different z-planes separated by 0.5 µm using a 63x oil-immersion objective (NA) at 2 frames per minute and using an exposure time of 100 ms and a 488 nm laser to image mGFP-PRC1 and 40 ms exposure time and a 405 nm laser imaging for Hoechst-stained. Fluorescence recovery after photobleaching (FRAP) experiments of mGFP-PRC1, Kif4A-mGFP and EGFP-CLASP1 were performed by bleaching a small area (0.6 µm diameter) in the microtubule overlap region using the 488 nm laser. Images were recorded in three different z planes separated by 0.5 µm at 30 frames per minute over a period of 3 min post bleaching using a 63x oil objective. Time -lapse imaging of mGFP-EB1 and mCherry-PRC1 co-expressing cells was performed in a single plane using a 100x oil immersion objective (NA) at 4 frames per second with exposure times of 50 ms (488 nm) and 30 ms (405 nm and 640 nm) over a period of 3 min.

### Determination of protein expression levels

Commercial antibodies were used to check the expression levels of proteins by western blot of cell lysate. EB1 and PRC1 proteins were detected by using antibodies sc-47704 (RRID:AB_2141631) and sc-376983 (both Santa Cruz Biotechnlogy), respectively. Tubulin was used as a loading control and detected using antibody ab18251 (RRID:AB_2210057) (Abcam). For detection of these primary antibodies, horse raddish peroxidase-coupled secondary antibodies were used.

### Average kymograph generation

Kymographs of metaphase, early anaphase, mid anaphase, late anaphase, and telophase cells expressing tagged endogenous EB1 (mGFP) and PRC1 (mCherry) were generated from movies imaged at 4 frames per second using the 100x objective. Kymographs of metaphase cells ectopically expressing EB1-mGFP (Suppl. Fig. 3F) were generated from movies imaged at 4 frames per second using the 100x objective. For each cell of interest, a manual tracking procedure was used to align the cell: for each frame the position of each centrosome was marked manually; the image was then translated and rotated to centre the midzone and align the spindle with the horizontal image axis. A kymograph was then automatically generated along the centrosomal axis averaging over a line width of 8 µm. Kymographs generated from individual cells were averaged together: for each kymograph the space (x) axis was normalized to the average metaphase centrosome distance, and the time (y) axis was shifted to align the start of anaphase, based on the DNA channel. For each fluorescence channel, a fluorescence profile of the average intensity in the midzone region was produced from the average kymograph for further analysis.

### Average fluorescence kymograph analysis

To extract the time course of the total PRC1 profile width (a measure for spindle length), the central PRC1 peak width (a measure of antiparallel overlap length), and the distance between separating chromosomes for Fig. 2D, the mathematical functions were used for fits to background-subtracted and bleaching-corrected intensity profiles (using OriginLab): A single Gaussian distribution was found to fit well metaphase PRC1 profiles, allowing to quantify its full width at half maximum (FWHM), i.e. the metaphase spindle length as labelled by PRC1. A sum of one central Gaussian peak with two equidistant side Gaussian peaks with identical heights and widths was used to fit anaphase PRC1 profiles; this allowed to quantify (i) the width of the entire profile (defined as the distance between the side peaks plus one FWHM of the side peaks) and (ii) the FWHM of the central PRC1 peak (could be determined reliably from t = 3 min onwards). A sum of 2 spatially displaced and otherwise identical Gaussian distributions was fit to the anaphase DNA profiles to extract the average distance between separating chromosomes. The maximum peak intensity was extracted from the PRC1 profiles by averaging the neighbouring 5 intensity values around the maximum value in each profile.

### FRAP analysis

Images were corrected for drift and centered by manually tracking centrosomes, as described above. Before FRAP, a 20 μm-long, 80 pixel-wide (8 μm) rectangular selection was drawn along the centrosomal axis to inspect the PRC1 and DNA distributions. The corresponding intensity profiles shown in Figure 3B (top) are background subtracted and have been normalized dividing them by their maximum intensity value. In order to analyse protein recovery after photobleaching, the fluorescence intensity was measured at the FRAP point (using a ROI of 0.6 µm radius circle) at every time point and normalized to the average intensity of the last 4 time points before FRAP. FRAP intensity curves from individual movies were classified into metaphase to telophase stage using the estimated time since mitosis and further refined using the PRC1and DNA intensity profiles. All FRAP intensity curves for a specific phase were averaged together and fit with the function *I* = *I*_*0*_ + *I*_a_(1-e^-t/τ^) (Fig. 3B, bottom) to obtain the recovery time τ, and the recovery fraction *m*_f_ *= I*_a_ /(1-*I*_*0*_) (Fig. 3C). In the case of KIFA-mGFP and EGFP-CLASP1, FRAP curves were also corrected for photobleaching by dividing them by the fluorescence intensity measured within a reference region.

### Time stack to visualize EB1 end tracking in interphase cells

The colour-coded time-stack images in Fig. 1A and Suppl. Fig.3E were generated using a previously developed ImageJ macro ^72^. Each individual frame of 20 time-lapse images acquired at 1s interval were colour-coded from red to blue sequentially in accordance to the rainbow order. Subsequently, the colour-coded time-lapse images were stacked using the z-stack option in ImageJ.

### Determination of EB1 velocity in different phases of mitosis

To quantify EB1 comet velocity, we generated kymographs from movies taken from cells in different phases of mitosis. The cells were imaged at 4 frames per second using a 100x oil objective and exposure times of 50 ms (488 nm) and 30 ms (405 nm and 640 nm). Kymographs were generated using FIJI drawing a line, with a width of 0.3µm, along the centrosomal axis and using the FIJI “Multi kymograph” function. Kymographs have been subsequently tracked using the web tool ‘KymoButler’ (Version 1.0.2; http://kymobutler.deepmirror.ai), which is based on techniques described by Jakobs *et al*.^73^ and relies on artificial intelligence to infer and reconstruct the trajectories. The parameter values used as inputs for tracking were ‘Threshold for track detection’ = 0.075, ‘Minimum size of detected objects’ = 3 pixel, and ‘Minimum number of consecutive frames’ = 3. The obtained trajectory dataset has then been manually polished by removing spurious trajectories and by reconnecting together those traces that had been split into different trajectories. Comets from the polished datasets were classified into “external” comets (when EB1was tracking astral microtubules) and “internal” comets (EB1 comet was moving in between of the two poles). EB1 comets velocity was finally calculated dividing the spatial length of each trajectory by its temporal duration, in order to obtain the average comet velocity.

## ACKNOWLEDGEMENTS

We thank the Flow Cytometry, the Cell Services and the Advanced Light Microscopy science technology platforms of the Francis Crick Institute for cell sorting, for providing wild-type hTERT-RPE1 and 293FT cells, and for help with some of the confocal microscopy imaging, respectively. We thank Anna Akhmanova, Stephen P. Jackson, Tim Mitchison, and Jennifer Tirnauer and for providing plasmids. We thank Julian Gannon for help with cell culture.

## AUTHOR CONTRIBUTIONS

**J.A**.: Conceptualization, cell lines development, experiments, data analysis, writing of the manuscript, figure preparation. **N.I.C**.: Microscopy support, data analysis, figure preparation. **D.N**. Microscopy support, data analysis, figure preparation. **W.M.L**.: Help with cell line development and figure preparation. **T.S**.: Conceptualization, data analysis, writing of the manuscript, supervision, funding.

