## Supplementary legends, movie legends and Supplementary figures for "Gradual Compaction of the Central Spindle Decreases its Dynamicity as Revealed in PRC1 and EB1 Gene-Edited Human Cells"

#### SUPPLEMENTARY FIGURE LEGENDS

**Supplementary Figure 1. EB1 localization in different cell lines. (A)** Fluorescence microscopy of examples of CRISPR/Cas9 modified RPE1 cells expressing both mGFP-EB1 (green) and mCherry-PRC1 (magenta) from their endogenous loci, showing co-localization of EB1 with PRC1 during different phases of mitosis. This behaviour of EB1 was considered unnatural, and therefore these cells were excluded from further study. **(B) - (D)** Control experiments: **(B)** Immunofluorescence microscopy of EB1 in unmodified RPE1 cells. **(C)** Immunofluorescence microscopy of EB1 in gene-edited RPE1 cells expressing only mGFP-PRC1 from its endogenous locus. **(D)** Immunofluorescence microscopy of EB1 in gene-edited RPE1 cells expressing only mCherry-PRC1 from its endogenous locus. The experiments in (B) - (D) demonstrate that endogenous, unmodified EB1 displays the typical "comet"-like localization to microtubule ends in RPE1 cells. DNA is always stained by Hoechst and shown in blue.

**Supplementary Figure 2. Characterization of mGFP-EB1 and mCherry-PRC1 expressing hTERT-RPE1 cells.**
**(A)** PCR test for correct mGFP gene insertion at the genomic EB1 gene locus. The schematic (top) indicates where the primers (small arrows) bind to the genomic DNA (black arrow head indicates the position of primer binding). One primer binds inside the mGFP gene, the other primer outside the left homology arm (LHA). Agarose gel (bottom) shows DNA fragments amplified from genomic DNA of control (Ctrl) and gene-edited single cell clones (clones A1-A5). The detection of a 2 kb band for the single cell clones demonstrates the correct insertion of the mGFP gene at the 5' end of the endogenous EB1 genes. (**B)** PCR test for determining the number of modified EB1 alleles. The schematic (top) indicates where the primers bind to the genomic DNA for this PCR test. One primer binds to the region outside the left homology arm (LHA) and the other primer binds inside the right homology arm (RHA). Agarose gel (bottom) showing DNA fragments amplified from control and gene-edited cells. The absence of the 1.2 kb band and the presence of a 2 kb band for the single cell clones A1-5 demonstrates that both EB1 alleles are modified in the gene-edited cells. **(C)** Western blot of cell lysates using an anti-EB1 antibody (top) showing the expression levels of unmodified EB1 in control cells and mGFP-EB1 in the gene-edited single cell clones A1-5 (left). Densitometry of the bands demonstrates that clones A1, A2, A3, A4 and A5 express 0.47, 0.45, 0.60, 0.33 and 0.0.27 times the amount of mGFP-EB1, respectively, compared to the amount of EB1 in unmodified control cells (n=2), data is normalized with respect to the tubulin control. Western blot of the same samples using an anti-tubulin antibody (bottom) as a loading control. Single cell clone B3, expressing mGFP-EB1 from its endogenous locus was selected for further gene-editing to additionally tag endogenous PRC1 with mCherry. **(D-F)** PCR tests and western blots, as indicated, for cells expressing both mCherry-PRC1 and mGFP-EB1 from their endogenous loci. All gene-edited clones (B1and B2) show apparently correct mCherry

gene insertion at the 5' end of the PRC1 gene (D) and have both PRC1 alleles tagged (E). Both clones B1 and B2 show correct protein expression (F). Clones B1, and B2, show expression levels of mCherry-PRC1 are 0.60, and 0.3 times of the expression levels of PRC1 in only EB1 modified control cells and unmodified control cells (n=2). Clone B1 was selected for further studies.

**Supplementary Figure 3. hTERT-RPE1 cells coexpressing endogenous mCherry-PRC1 and ectopic EB1-GFP.**

(**A-C)** PCR tests and western blots, as indicated, for gene-edited cells expressing mCherry-PRC1 from the endogenous locus. Assays as described in Suppl. Figs. 2. All gene-edited clones (C1-4) show correct mCherry gene insertion at the 5' end of the PRC1 gene (A) and have both PRC1 alleles tagged (B). Clones C1, C2, C3, and C4, express 0.32, 0.51, 0.87, and 0.41 times the amount of mCherry-PRC1, respectively, compared to the amount of PRC1 in unmodified control cells (n=2). Clone C3 was selected for lentiviral infections to randomly insert an ectopic EB1-GFP gene. **(D)** Western blot of cell lysates using an anti-EB1 antibody (left) showing expression levels of ectopically expressed EB1-eGFP from a randomly integrated gene after lentiviral transfection of cells also expressing mCherry-PRC1 and unmodified EB1 from their endogenous loci. Clone D1 shows expression of ectopic EB1-GFP and unmodified endogenous EB1. EB1-GFP expresses 3.4 fold more than unmodified EB1 in unmodified control cells (n=2). Western blot of the same samples using an anti-tubulin antibody (right) as a loading control. **(E)** Spinning disk confocal microscopy image of an interphase cell (left) and time stack from a time lapse movie of the same cell (right) showing ectopically expressed EB1-GFP microtubule end tracking traces color-coded as a function of time. **(F)** Image of a metaphase cell (left) and kymograph along the spindle axis of the same cell (right) showing EB1-GFP tracks along the spindles as well as astral microtubules. **(G)** Average kymograph generated along the spindle pole-to-pole axis from 10 mitotic movies, showing Hoechst-stained DNA, EB1 with a C-terminal GFP tag and mCherry-PRC1. Scale bar = 10 μm.

**Supplementary Figure 4. EB1 comet characterization in late phases of mitosis.**

**(A**) Still image from a late anaphase movie showing how kymographs were generated along the centrosome axis by drawing a line from pole to pole (white line in figure). **(B)** EB1 tracks are shown for microtubules in different phases of mitosis. DNA is shown in blue. Pole positions are indicated by the vertical dashed black lines. **(C)** EB1 comet speed distributions in different phases of mitosis. EB1 comet speed decreases from metaphase to late anaphase. Box plots show the 25%-75% interval of the distributions, black squares indicate mean values, horizontal black lines show median values, and vertical lines indicate the 1%-99% interval of the distributions. **(D)** Survival probability of EB1 comet track length (expressed both in seconds and frames) for the different phases of mitosis. **(E)** Histograms of EB1 comet location along the pole-to-pole axis for the different mitosis stages. (C-E) The dataset of EB1 comets and cells considered in the analysis is the same than in Figure 1E.

**Supplementary Figure 5. Characterization of CRISPR/Cas9 gene-edited hTERT-RPE1 cells expressing mGFP-PRC1.**

**(A)** PCR test for correct mGFP gene insertion at the genomic PRC1 locus. The schematic (top) indicates where the primers (small arrows) bind to the genomic DNA for this PCR test (black arrow head indicates the position of primer binding). One primer binds to the region outside the right homology arm (RHA) and the other primer binds inside the mGFP gene (outside the right homology arm (RHA)). No band is expected for the control and for the modified allele a 2 kb band is expected. Agarose gel (bottom) showing DNA fragments amplified from genomic DNA of control (Ctrl) and gene-edited single cell clones (E1-5) by PCR. The detection of a 2 kb band only for the single cell clones demonstrates the correct insertion of the mGFP gene at the 5' end of the endogenous PRC1 gene in the gene-edited clones. **(B)** PCR test for determining the number of modified alleles. The schematic (top) indicates where the primers bind to the genomic DNA for this PCR test. One primer binds to the region outside the left homology arm (LHA) and the other primer binds inside the right homology arm (RHA). The expected band size for the control is ~1.2 kb and for the modified allele is ~2 kb. Agarose gel (bottom) showing DNA fragments amplified from genomic DNA of unmodified control cells (Ctrl) and gene-edited single cell clones (E1-5) by PCR. The absence of the 1.2 kb band and the presence of a 2 kb band for the single cell clones demonstrates that both PRC1 alleles are modified in the single cell clones tested. **(C)** Western blot of cell lysate using an anti-PRC1 antibody (left) showing the expression levels of unmodified PRC1 in control cells and mGFP-tagged PRC1 in the gene-edited single cell clones. Double bands indicate that at least two isoforms of PRC1 are expressed both in control and gene-edited cells. Densitometry of the bands demonstrated that clones E1, E2, E3, E4 and E5 show expression levels of mGFP-PRC1 that are 0.96 ± 0.15, 0.97 ± 0.13, 0.94 ± 0.14, 0.0.92 ± 0.14 and 0.93 ± 0.14 times of the expression levels of PCR1 in unmodified control cells (n=5). Clone E1 was selected for further studies. A western blot of the same samples using an anti-tubulin antibody (right) was made as a loading control.

**Supplementary Figure 6. Average distribution of mGFP-PRC1 in the mitotic spindle at different time points during mitosis.**

**(A)** Separate mGFP-PRC1 (left) and Hoechst-stained DNA (right) channels of the average kymograph from Fig. 2B right panel showing the PRC1 and DNA distributions along the spindle axis throughout mitosis. **(B)** Individual average PRC1 profiles (black lines) for different time points (-3, 0, 1, 3, 7 and 10 min). The shadow over the profile line represents the standard error. (C) Comparison of the fluorescence recovery curves from (Fig. 3B bottom) here in a single graph. Error bars indicate the standard error of the mean.

#### SUPPLEMENTARY MOVIE LEGENDS

**Supplementary Movie S1:** Spinning disk confocal microscopy live-cell imaging of hTERT-RPE1 cells expressing endogenous mGFP-EB1, showing microtubule plus end tracking of mGFP-EB1 in interphase and anaphase cells. EB1 tracks interphase, astral, and mitotic microtubules. Interphase images were captured at 4 frames/s, anaphase images were captured at 30 frames/min, and are displayed at 10 frames/s The time stamp is min:s.

**Supplementary Movie S2**: Spinning disk confocal microscopy live-cell imaging of hTERT-RPE1 cells coexpressing endogenous N-terminally mGFP-tagged EB1 (green) and endogenous mCherry-PRC1 (magenta), showing the localization of both fluorescently tagged protein in mitosis. EB1 tracks astral, and mitotic microtubules. Chromosomes (blue) were stained with Hoechst 33342. Images were captured at 12 frames/min, and are displayed at 10 frames/s. The time stamp is min:s.

**Supplementary Movie S3**: Spinning disk confocal microscopy live-cell imaging of hTERT-RPE1 cells coexpressing endogenous N-terminally mGFP-tagged EB1 (green) and endogenous mCherry-PRC1 (magenta), showing the localization of both fluorescently tagged protein in metaphase, early anaphase, mid anaphase, late anaphase and telophase. EB1 tracks astral, and mitotic microtubules. Chromosomes (blue) were stained with Hoechst 33342. Images were captured at 4 frames/s, and are displayed at 10 frames/s. The time stamp is min:s.

**Supplementary Movie S4**: Spinning disk confocal microscopy live-cell imaging of hTERT-RPE1 cells expressing endogenous mGFP-PRC1. PRC1 localizes to spindle microtubules after nuclear envelope breakdown, binds to microtubule bundles in the metaphase spindle, and then accumulates at antiparallel microtubule overlaps in the midzone of anaphase spindles and in the midbody in telophase. Chromosomes (blue) were stained with Hoechst 33342. Images were captured at 2 frames/min, and are displayed at 10 frames/s. The time stamp is min:s.

**Supplementary Movie S5**: Spinning disk confocal microscopy live-cell imaging of hTERT-RPE1 cells expressing endogenous mGFP-PRC1, showing fluorescence recovery after photobleaching (FRAP) of mGFP-PRC1 in different phases of mitosis. The recovery of mGFP-PRC1 slows down with the progression of mitosis. Images were captured at 30 frames/min, and are displayed at 3 frames/s. The time stamp is min:s.

**Supplementary Movie S6**: Spinning disk confocal microscopy live-cell imaging of hTERT-RPE1 cells stably expressing ectopic KIF4A-mGFP, showing fluorescence recovery after photobleaching (FRAP) of KIF4A-mGFP in late anaphase and telophase. The recovery of KIF4A-mGFP slows down from late anaphase and telophase. Images were captured at 30 frames/min, and are displayed at 3 frames/s. The time stamp is min:s.

**Supplementary Table 1. Summary of cell lines used in this study.**

| hTERT-RPE1 cell lines | Clone number | Main Figures | Suppl. Figures | Suppl. Movies |
|---|---|---|---|---|
| mGFP-EB1 (endogenous) | A3 | Figure 1 | Suppl. Figure 2 | Suppl. Movie 1 |
| mGFP-EB1 (endogenous) & mCherry-PRC1 (endogenous) | B1 | Figure 1 | Suppl. Figure 2, 4 | Suppl. Movie 2, 3 |
| mCherry-PRC1 (endogenous) | C3 | | Suppl. Figure 3 | |
| mCherry-PRC1 (endogenous) & EB1-GFP (ectopic) | D1 | | Suppl. Figure 3 | |
| mGFP-PRC1 (endogenous) | E1 | Figure 2, 3 | Suppl. Figure 5, 6 | Suppl. Movie 4, 5 |
| mCherry-PRC1 (endogenous) & KIF4A-mGFP (ectopic) | | Figure 4 | | Suppl. Movie 6 |
| mCherry-PRC1 (endogenous) & EGFP-CLASP1 (ectopic) | | Figure 5 | | |
| unmodified control | | | Suppl. Figure 1, 2, 3, 5 | |

### Supplementary Figure 1

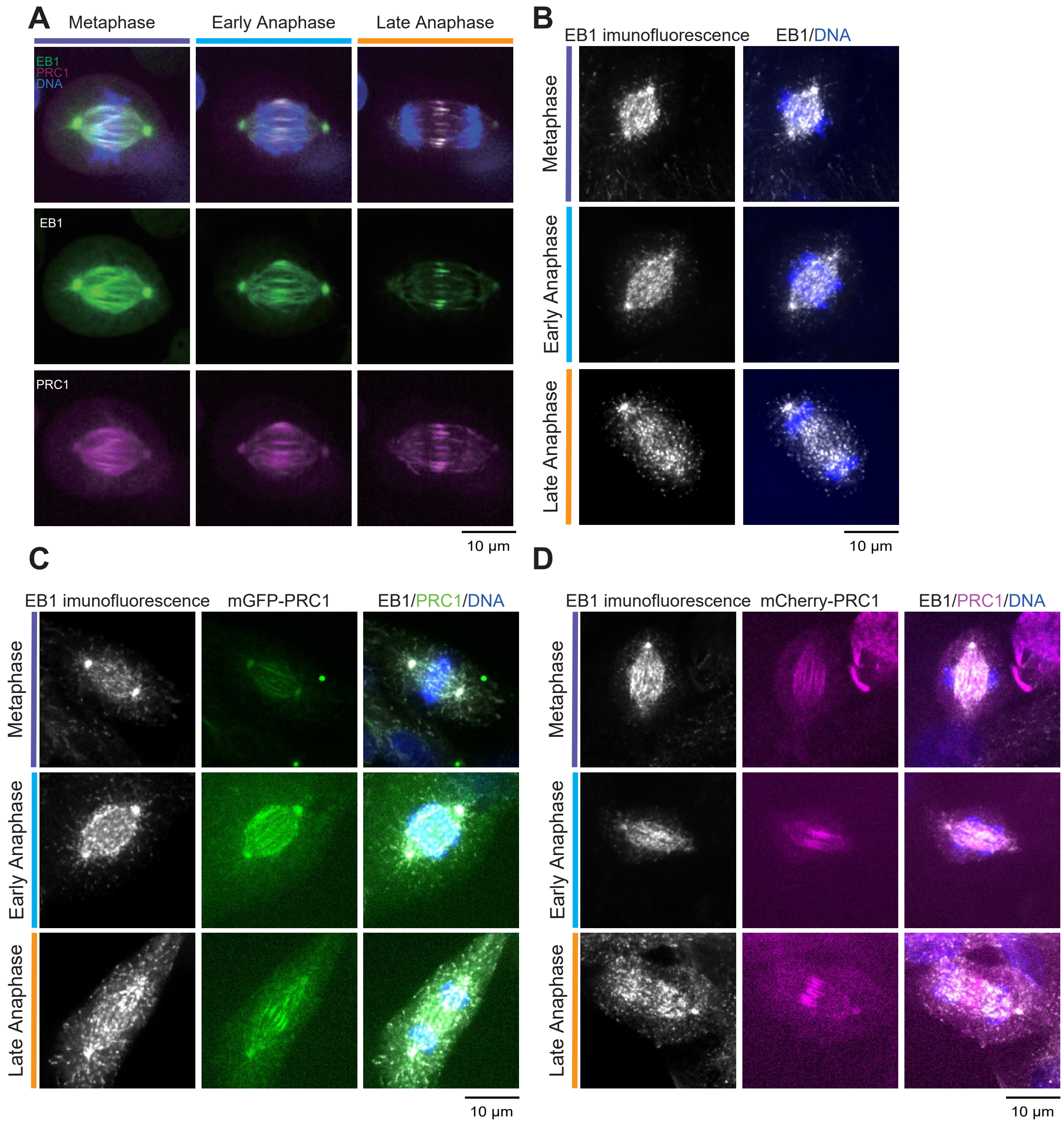

### Supplementary Figure 2

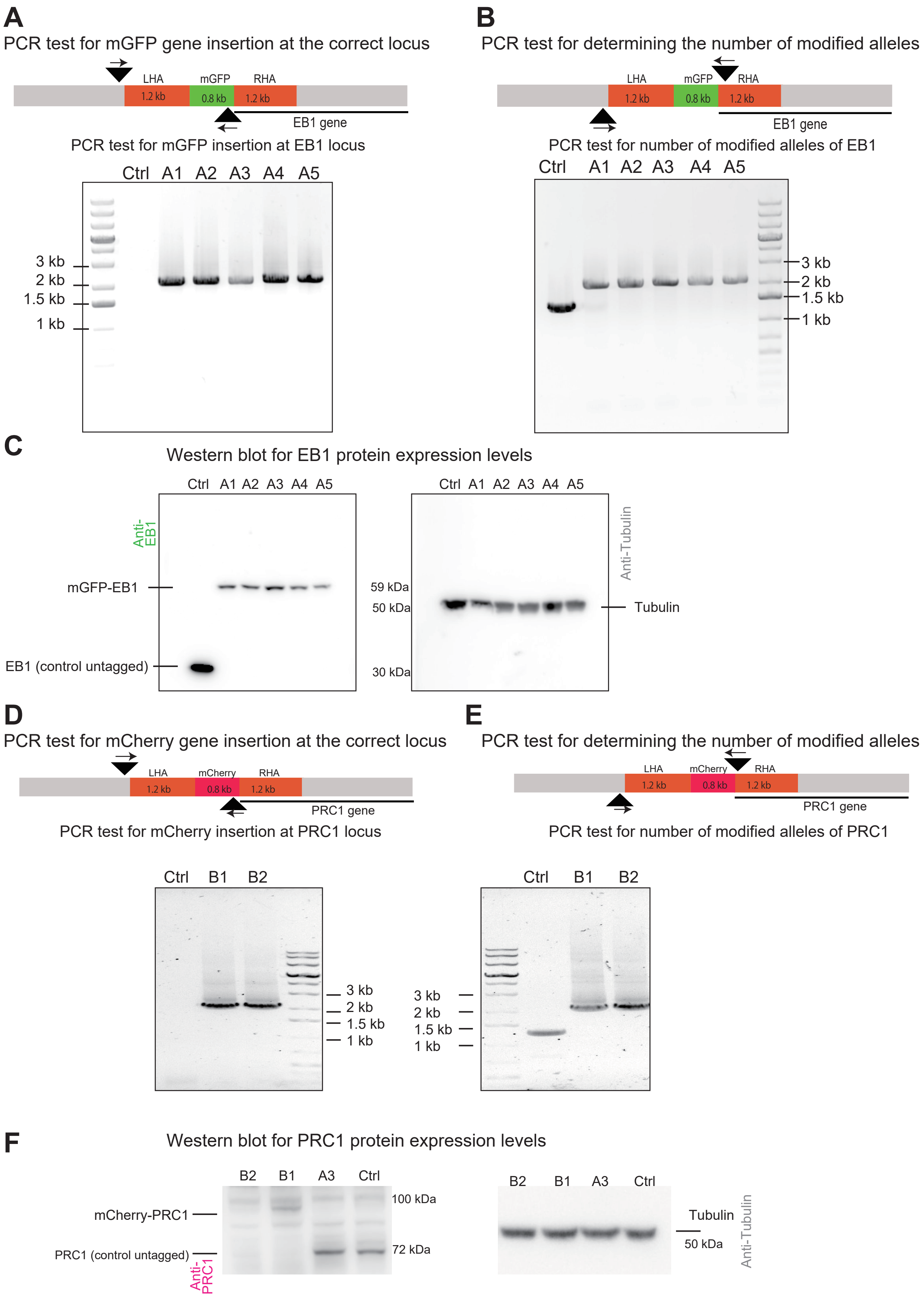

### Supplementary Figure 3

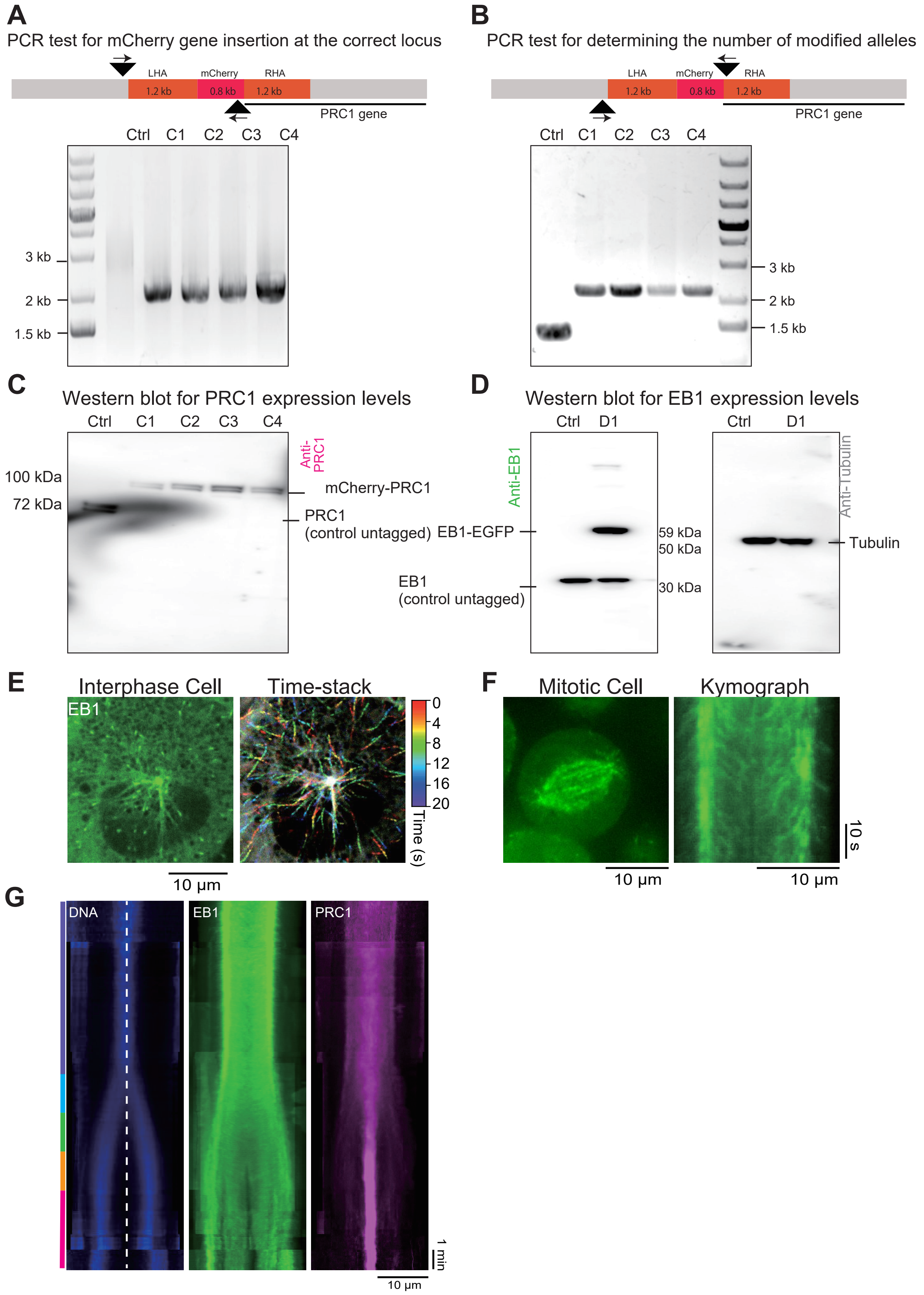

### Supplementary Figure 4

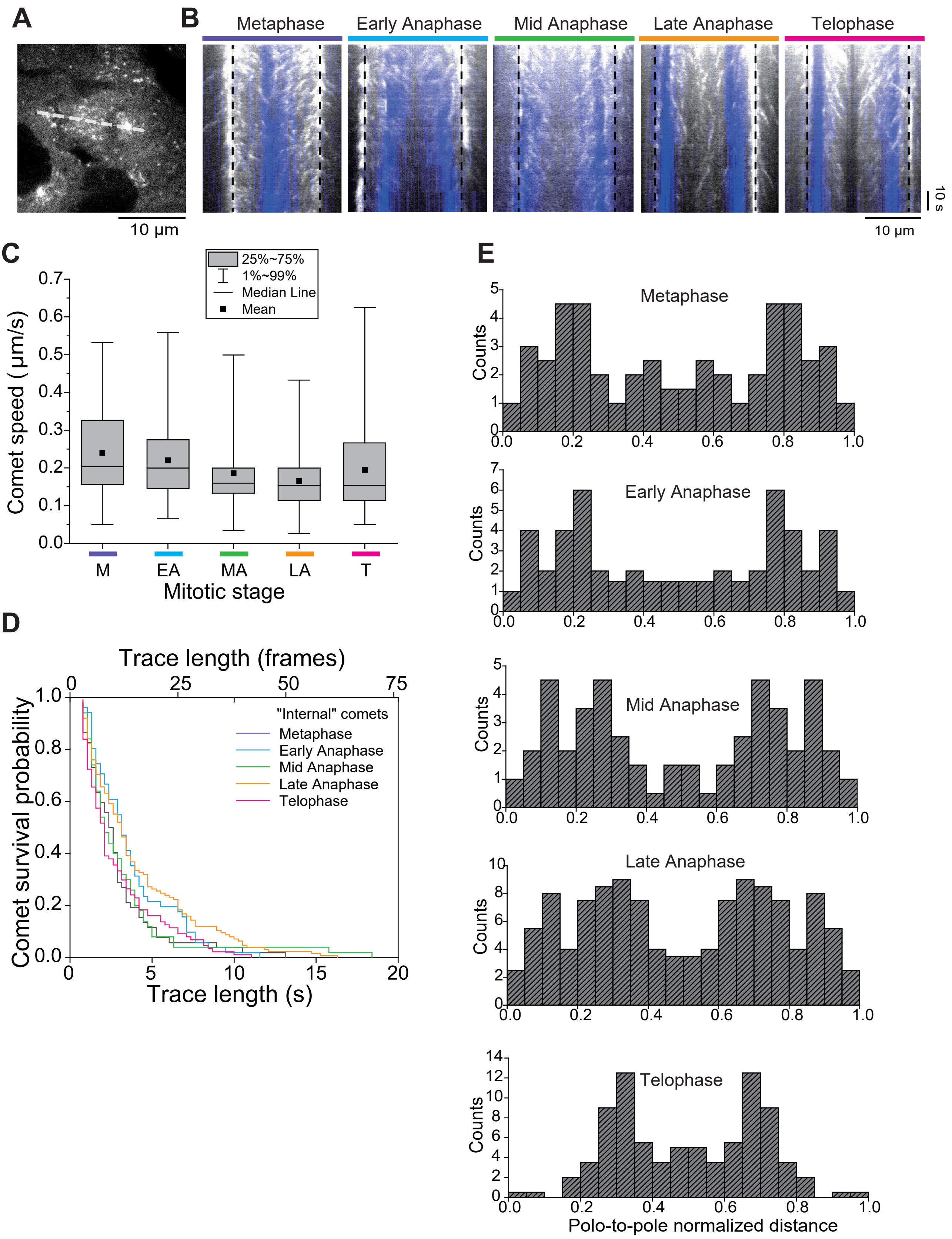

### Supplementary Figure 5

**A**

PCR test for mGFP gene insertion at the correct locus

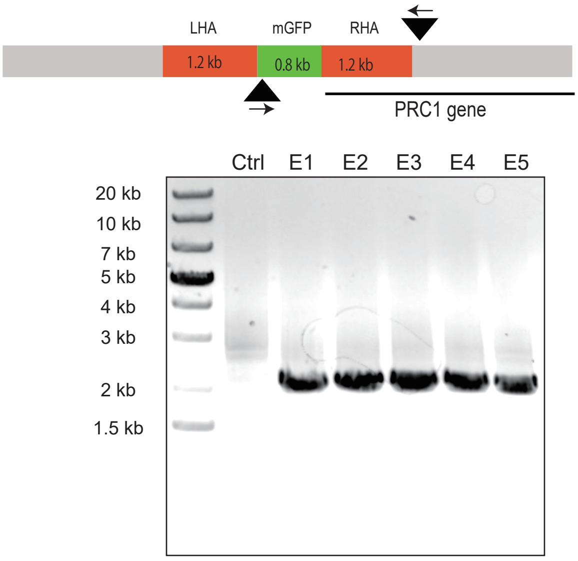


**B**

PCR test for determining the number of modified alleles

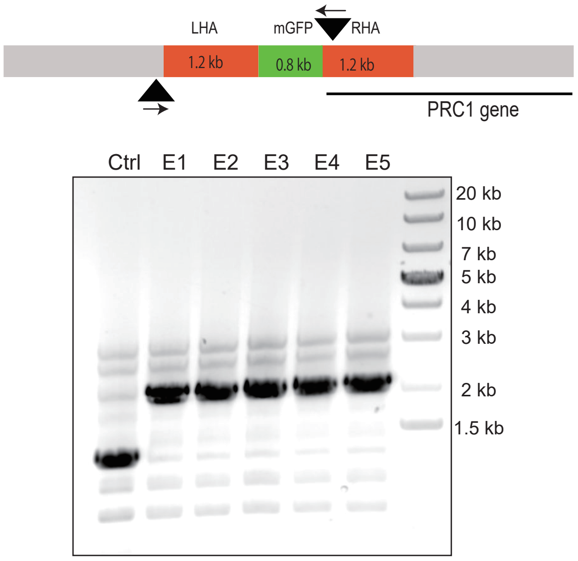


**C** Western blot for PRC1 protein expression levels

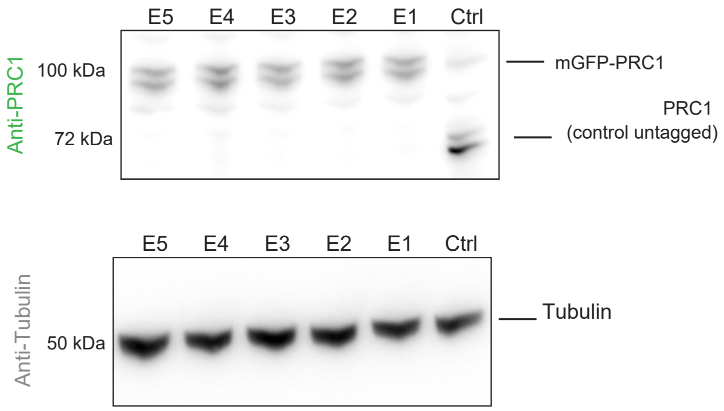

### Supplementary Figure 6

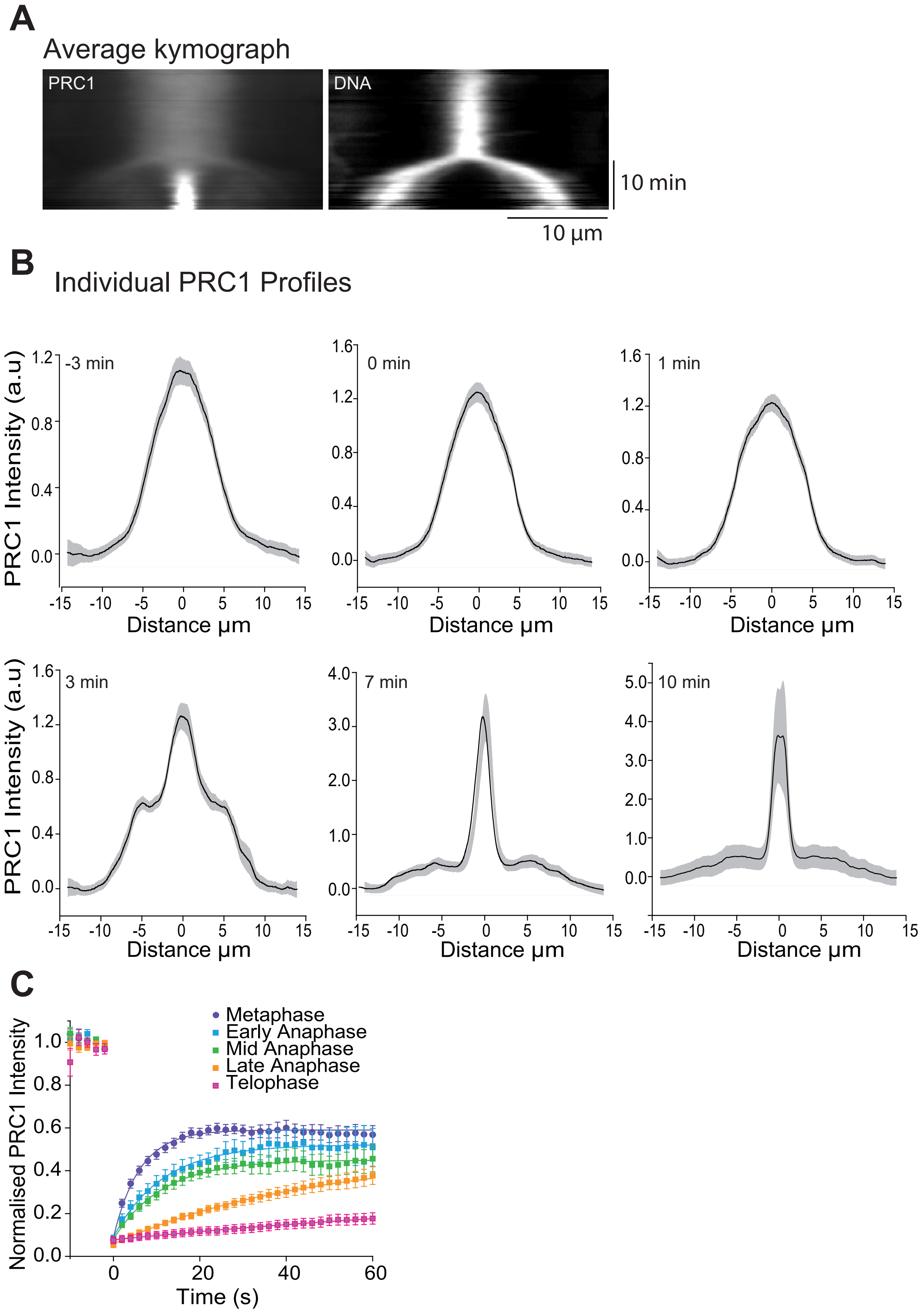

#### PRC1 donor vector primers

| | |
|---|---|
| LHA(PRC1-N)FP | TCGGTACCCGGGGATCGAGCAAAGATGTTTAGAAGTACAATAG |
| LHA(PRC1-N)RPGFP | CCTTGCTCACGCTCATGGCGGACGCTCCAAGCAG |
| LHA(PRC1-N) RPmcherry | CGCCCTTGCTCACCATGGCGGACGCTCCAAGCAG |
| mGFP (PRC1-N)FP | ATGAGCGTGAGCAAGGGCGA |
| mGFP(PRC1-N)RP | CGCACCTTCTCCTCATTCCGCCTCCTCCGCCCGCGTATCCGCCTCCCTTG |
| Mcherry(PRC1-N)FP | ATGGTGAGCAAGGGCGAGGA |
| Mcherry(PRC1-N)RP | CGCACCTTCTCCTCATTCCGCCTCCTCCGCCCTTGTACAGCTCGTCCATGCC |
| RHA(PRC1-N)FP | ATGAGGAGAAGGTGCGGGTTG |
| RHA(PRC1-N)RP | TCGACTCTAGAGGATCTTGAGACGTAGTCTCACTCTGTC |

#### SDM primers for the PAM motifs

| | |
|---|---|
| SDMguideA FP | TTGTTGCTCTCGGGGGGGTGTGGAGTAGGTCT |
| SDMguideA RP | AGACCTACTCCACACCCCCCCGAGAGCAACAA |
| SDMguideB FP | AGCGTCCGCCATGAGAAGAAGGTGCGGGTT |
| SDMguideB RP | AACCCGCACCTTCTTCTCATGGCGGACGCT |
| SDMguideC FP | GTGGAGTAGGTCTGGAGGTGGACTCACGGCTGCTT |
| SDMguideC RP | AAGCAGCCGTGAGTCCACCTCCAGACCTACTCCAC |
| SDMguideD FP | CGGGGAAATCGACCGGACGCGGGAG |
| SDMguideD RP | CTCCCGCGTCCGGTCGATTTCCCCG |

#### Guide RNA sequence

| | |
|---|---|
| Guide A1 | ACCGAGGTCCAGACCTACTCCACA |
| Guide A2 | AAACTGTGGAGTAGGTCTGGACCT |
| Guide B1 | ACCGTGCTTGGAGCGTCCGCCATG |
| Guide B2 | AAACCATGGCGGACGCTCCAAGCA |
| Guide C1 | ACCGCTCCAAGCAGCCGTGAGTCC |
| Guide C2 | AAACGGACTCACGGCTGCTTGGAG |
| Guide D1 | ACCGTGCGGGTTGCGGGGAAATCG |
| Guide D2 | AAACCGATTTCCCCGCAACCCGCA |

#### Analysis Primers

| | |
|---|---|
| FPPRC1N-Terminus OutSide LH/ | TGCCAAACAAGGAAATGCCAGTAT |
| RPPRC1N-Terminus inside RHA: | CAACCCGCACCTTCTCCTCA |
| RPPRC1N-Terminus Outside RH. | CCTTGATAGAGGAGATGCCTAC |
| mGFP (PRC1-N)FP | ATGAGCGTGAGCAAGGGCGA |
| mCherry(PRC1-N)FP | ATGGTGAGCAAGGGCGAGGA |
| mCherry(PRC1-N)RP | CGCACCTTCTCCTCATTCCGCCTCCTCCGCCCTTGTACAGCTCGTCCATGCC |

#### Sequencing Primers

| | |
|---|---|
| mcherryRPseqforPRC1 | TGAACTCCTTGATGATGGCCATGT |
| PRC1 LHAFPseq | GTCACGGGCATGCGTGCACA |
| mGFP primer | GTCCTTAAGGAGTTCGTGACCG |

#### EB1 donor vector primers

| | |
|---|---|
| EB1 FP left H arm | TCGGTACCCGGGGATCGGAAATAGGATCTCACTGCC |
| EB1 RP left H arm | CCTTGCTCACGCTCATCTTCTAAAGCATGGGAAGAAAAG |
| mGFP FP EB1 | ATGAGCGTGAGCAAGGGCGA |
| mGFP RP EB1 | TATACGTTCACTGCCATGCCATCCACCGCGCCTTCGCCGCCGCTGCCCGC GCCGCCGCTGCCGCCCGCCTGCGCCGCGTATCCGCCTCCCTTG |
| EB1 FP Right H arm | ATGGCAGTGAACGTATACTCAA |
| EB1 RP Right H arm | TCGACTCTAGAGGATCACCACACCGAGACTTTAAATCA |

#### SDM primers for the PAM motifs

| | |
|---|---|
| EB1 SDM guide A FP | GAACAGTTGTGCTCAGTTAAGAGAAATCTGCTG |
| EB1 SDM guide A RP | CAGCAGATTTCTCTTAACTGAGCACAACTGTTC |
| EB1 SDM guide B FP | GACATGACATGCTGGCTTGGATCAATGAGTCTC |
| EB1SDM guide B RP | GAGACTCATTGATCCAAGCCAGCATGTCATGTC |
| EB1 SDM guide C FP | GACATGACATGCTGGCTTGGATCAATGAGTCTC |
| EB1SDM guide C RP | GAGACTCATTGATCCAAGCCAGCATGTCATGTC |
| EB1 SDM guide D FP | CAGGTAAGAGAAATCTGCTTTATCATTTTTCTAGGAAAGC |
| EB1 SDM guide D RP | GGCTTTCCTAGAAAAATGATAAAGCAGATTTCTCTTACCTG |

#### Guide RNA sequence

| | |
|---|---|
| Guide A1 | ACCGAAGATCGAACAGTTGTGCTC |
| Guide A2 | AAACGAGCACAACTGTTCGATCTT |
| Guide B1 | ACCGCTGCAGAGACTCATTGATCC |
| Guide B2 | AAACGGATCAATGAGTCTCTGCAG |
| Guide C1 | ACCGCTGCAGAGACTCATTGATCC |
| Guide C2 | AAACGGATCAATGAGTCTCTGCAG |
| Guide D1 | ACCGCTCAGGTAAGAGAAATCTGC |
| Guide D2 | AAAC GCAGATTTCTCTTACCTGAG |

#### Analysis Primers

| | |
|---|---|
| EB1FPoutsideLHA | CTGTAAGGTCATTTGATACTGCC |
| EB1RPinsideRHA | GTCATGTCGACTTAGGTTATCAC |
| mGFP FP EB1 | ATGAGCGTGAGCAAGGGCGA |
| mGFP RP EB1 | TATACGTTCACTGCCATGCCATCCACCGCGCCTTCGCCGCCGCTGCCCGC GCCGCCGCTGCCGCCCGCCTGCGCCGCGTATCCGCCTCCCTTG |

#### Sequencing primers

| | |
|---|---|
| mGFPseq for EB | GTCCTTAAGGAGTTCGTGACCG |
| EB1 RHA reverse | CTCTGTGTGTGGCTTTGCAGTC |

#### EB1-GFP Lenti viral Cloning Primers

| | |
|---|---|
| Forward Primer | CCGCTCGAGATGGCAGTGAACGTATACTCA |
| Reverse Primer | CGCGGATCCCGCGTATCCGCCTCCCTTG |

#### KIF4A-mGFP Lenti viral Cloning Primers

| | |
|---|---|
| Forward Primer | CTCAAGCTTCGAATTATGAAGGAAGAGGTGAAGGGAAT |
| Reverse Primer | TAGAGTCGCGGGATCTTACTTGTACAGCTCGTCCATG |

#### EGFP-CLASP1 Lenti viral Cloning Primers

| | |
|---|---|
| Forward Primer | CTCAAGCTTCGAATTATGGTGAGCAAGGGCGAGGA |
| Reverse Primer | TAGAGTCGCGGGATCTTAGCTGTGCGTGGAGACATC |
